# Genome and transcriptome mining revealed evolutionary insights and tissue-specific expression patterns of Cytochrome P450 superfamily in *Aquilaria sinensis*

**DOI:** 10.1101/2023.08.04.551930

**Authors:** Ankur Das, Khaleda Begum, Raja Ahmed, Suraiya Akhtar, Sofia Banu

## Abstract

The Cytochrome P450 (CYPs) enzyme superfamily has evolved and expanded in plants to play a significant role in the biosynthesis of valuable secondary metabolites. In *Aquilaria* plant, the process of wounding and fungal infection results in the accumulation of distinct aromatic metabolites which contribute to the formation of resinous agarwood. The members of CYPs in plants genomes have diversified to catalyze a wide array of fragrant metabolites. Therefore, this study aimed to identify and provide insights into the evolution and expansion of the CYP superfamily members in *Aquilaria sinensis* and elucidate their tissue-specific functional role through mapping and expression analyses. In total, 179 AsCYPs were identified and subsequently classified into 8 clans and 42 families which were found to be dispersed in the 8 chromosomes. Duplication analysis highlights slow segmental events as a major force, coupled with negative selection pressure, behind the expansion of the *AsCYPs*. We observed their participation in the biosynthesis of various secondary metabolites, particularly sesquiterpenoids. Expression analysis showed variation in the expression pattern of the genes in different tissues, revealing that the diversification of the AsCYP superfamily occurred to carry out tissue-specific functional role. Additionally, molecular docking of a sesquiterpene oxidase which is specifically expressed in wounded wood, indicated its potential to generate sesquiterpenoids derivatives in agarwood. This study sheds light on the evolution and expansion of the *AsCYPs* in the genome of *A. sinensis* and highlights their crucial role in the biosynthesis of various secondary metabolites found in different parts of the plant. Further functional exploration may pave the way for advancements in the field of *Aquilaria*-based fragrance development and natural product synthesis.

## Introduction

*Aquilaria sinensis* is a valuable plant species noted for producing a unique and fragrant resinous wood called agarwood. It has been utilized in medicine, perfumes, and religious rites for decades (López-Sampson and Page 2018). But the rising demand has resulted in the overexploitation of natural *Aquilaria* populations. (Hashim et al. 2016). The distinctive aroma of agarwood is attributed to the presence of specific fragrant metabolites such as sesquiterpenoids and phenyl chromones, which the tree synthesizes only after wounding and infection (Gao et al. 2019). In addition, the species owes its medicinal value to the metabolites, such as flavonoids, lignans, diterpenoids, and xanthones, which are found in their bark, stem, leaves, and flowers (Kristanti et al. 2018, Li et al. 2020). These properties have fueled the popularity of agarwood and *Aquilaria* plants in perfumery and pharmaceutical industries. The Cytochrome P450 (CYPs) superfamily holds a significant role as a key player in the biosynthesis and diversification of secondary metabolites in plants (Banerjee et al. 2018). It also actively participates in the processes associated with hormone signaling (Xu et al. 2015). The superfamily is composed of multiple-family (clan 71, clan 72, clan 85, and clan 86) and single-family clans (clan 51, clan 74, clan 97, clan 710, clan 711, and clan 727) (Nelson et al. 2004, Nelson et al. 1996). The members of clan 71 are grouped into A-type, while the rest members into non-A type (Paquette et al. 2000, Nelson 1999). Moreover, transcriptomics and expression based study carried out in certain plants have revealed their functional role during biotic stress caused by insects (Schuler et al. 1996) and pests (Li et al. 2010), as well as abiotic stresses such as drought (Tamiru et al. 2015), temperature (Qin et al. 2008), salinity (Wang et al. 2017), and heavy metals (Rai et al. 2015).

Certain members within the Cytochrome P450 (CYPs) superfamily perform essential and consistent functions in various pathways. For instance, the 51 clan is involved in sterol biosynthesis, the 97 clan in carotenoid biosynthesis, the 74 clan in oxylipid biosynthesis, and the 72 clan in hormone homeostasis. The members of 71 clans, however, actively participate in multiple pathways related to phenylpropanoids, terpenoids, and flavonoids biosynthesis (Hansen et al. 2021). These versatile enzymes have been the subject of extensive studies within various plant species, such as *Arabidopsis thaliana*, *Oryza sativa*, *Vitis vinifera*, and *Glycine max* (Nelson et al. 2004, Wei et al. 2018, Jiu et al. 2020, and Khatri et al. 2022). Through rigorous investigation, scientists have successfully identified a total of 245, 326, 236, and 317 *CYPs* in their respective genomes. Further, these genes have been meticulously categorized into different families and subfamilies. In plants, the CYP superfamily is considered as largest gene family, and owing to its complexity, substrate specificity and enzyme promiscuity, their expression pattern in tissues are highly variable and diverse (Awasthi et al. 2015, Hansen et al. 2021). In *O. sativa*, expression patterns of the members varied in different tissues, where few members exclusively expressed in particular tissue (Wei et al. 2018). Similarly, in *V. vinifera*, the CYP74B and CYP81D subfamily members consistently expressed in all the experimented tissue, while few highly expressed only in the leaves and buds. The expression of CYP71A subfamily was relatively low in all tissues (Jiu et al. 2020). The rapid evolution of CYPs have given plants the ability to adapt to their surroundings, and key factors driving the diversification and expansion include duplication and divergence events in the genome (Jaillon et al. 2007). In addition, duplication types including segmental, tandem, and rearrangements at the gene and chromosome levels contribute immensely to the evolution of protein families (Maher et al. 2006). Evidence supports segmental (*A. thaliana* and *G. max*) and tandem duplication (*V. vinifera*) events as major driving forces for CYP family expansion (Yu et al. 2017, Guttikonda et al. 2010, Jiu et al. 2020). Moreover, selection pressure on the duplicated members lead to neo/sub-functionalization of the proteins (Wanger et al. 2002)

In *Aquilaria* plants, although the metabolic pathways for biosynthesis of the metabolites are poorly understood till date. In our previous study, we attempted to systematically identify the *CYPs* of *Aquilaria agallocha* in a genome-wide level and comprehensively illustrated their functional role in secondary metabolites biosynthesis (Das et al. 2023). However, due to missing chromosome-level assembly of the genome and existence of the genome as numerous short scaffolds, limited number of duplicated genes were identified. Therefore, we could not emphasize and comprehend on the evolution and expansion of the superfamily. Additionally, sparse tissue-wide transcriptomic data of *A. agallocha* in NCBI Sequence Read Archive database and expression study being limited to one or two tissue type restricted our understanding. Nevertheless, recently, advancements in next-generation sequencing technologies and high-end genome assemblers have made it possible to sequence, assemble, and precisely annotate the genome of *Aquilaria sinensis* (Lour.) Spreng up to chromosomal-level (Ding et al. 2020). On top of that, the availability of transcriptomic data from different tissues, including leaf, bud, callus, aril, seed, stem, and wounded tree, has opened up new avenues for studying the expression and function of the genes. (Lv et al. 2022, Nong et al. 2020). Therefore, this study aims to utilize these resources, i.e., the genomic and transcriptomic data and the information available in the public database, to conduct genome-wide duplication and tissue-wide expression analyses of the *CYPs* in *A. sinensis*. Through this study, we aim to gain a better understanding on the effects of duplication and expansion of the family members, and provide valuable insights into the potential candidate members, their tissue-specific functions, and open possibilities for their functional validation and biotechnological and industrial exploration.

## 2. Methods and Materials

### 2.1. Identification and Phylogenetic Relation of CYP Superfamily in *Aquilaria sinensis*

The sequence data for *Aquilaria sinensis* (Lour.) Spreng was accessible through the BioProject accession number PRJNA556948 on NCBI. The P450 domain (PF00067) sequences specific to Viridiplantae were extracted from InterPro EMBL-EBI database. The hmmbuild command was used in HMMER 3.0 to generate a hidden Markov model based on these P450 domains which was subsequently used with the hmmscan module to identify the CYPs from the protein sequences of *A. sinensis* (Potter et al. 2018). Protein sequences lacking the P450 domains and conserved motifs, as confirmed by the ScanProsite tool (Sigrist et al. 2002) and MEME Suite (Liu et al. 2018), respectively were excluded from further analysis. The remaining sequences were then subjected to blast search against the Uniprot database using a threshold e-value of 1e-10 to confirm their identity. The molecular weights and theoretical pI of the proteins were calculated using ExPASy server. Once the candidate *A. sinensis* CYPs (AsCYPs) were identified, protein sequences of *A. thaliana* were retrieved from TAIR (https://www.arabidopsis.org/) for phylogenetic analysis. Multiple sequence alignment was performed using CLUSTALW with default parameters, and a tree was drawn utilizing Neighbor-Joining (NJ) method in the MEGA software, with 1000 bootstrap replication and the p-distance model (Kumar et al. 2018).

### 2.2. Chromosomal Localization, Collinearity, Duplication and Selection Type Analysis

To determine physical locations of the *AsCYPs* on the chromosome, genomic information was extracted from BioProject accession number PRJNA556948, and depicted using TBtools (Chen et al. 2020). The duplication analysis in the genome of *A. sinensis* were procured through protein blast, where >75% of sequence similarity and query coverage were utilized to ascertain the linked duplicated members. The loci of two linked members, if existed within a 100 kb distance on the single chromosome, were considered to have arisen from tandem duplication. However, if the distance exceeds 100 kb, whether on the same or different chromosomes, it was classified as segmental duplication. (Islam et al. 2019, Lopez-Ortiz et al. 2019). The syntenic blocks for collinearity analysis for determining the evolutionary related CYPs were acquired through MCScanX software from the genomic and general feature format (GFF) data of *A. sinensis* and *A. thaliana* (Wang et al. 2012). Diversion time were calculated from the Synonymous (Ks) and non-synonymous (Ka) rate of substitution utilizing KaKs_calculator (Aslam et al. 2021, Moniz de Sa and Drouin 1996).

### 2.3. KEGG mapping and Functional Annotation

GO and InterPro terms were assigned to the AsCYPs to elucidate their involvement in biological, molecular and cellular processes using the web servers (Ashburner et al. 2000, (Mitchell et al. 2019). For pathway mapping and enzyme annotation, the protein sequences were searched against the Kyoto Encyclopedia of Genes and Genomes (KEGG) database (http://www.kegg.jp/blastkoala/). Sequences with missing information in KEGG were manually annotated based on homology to biochemically characterized proteins of Uniprot database (https://www.uniprot.org/) with parameters set to sequence similarity and alignment length and threshold e-value of 1e-10.

### 2.4. Expression Pattern Analysis of *AsCYPs* in RNA-seq data

To analyze the tissue-wide expression patterns of the members, RNA-seq raw reads of wounded and control wood tissue of *Aquilaria sinensis* (Ordinary and Chi-Nan germplasm) (BioProject number PRJNA819559) (Lv et al. 2022); leaves, flower, buds, aril, seed, and wounded stem (BioProject number PRJNA534170) (Nong et al. 2020); and salt treated callus tissue (BioProject number PRJNA305967) (Wang et al. 2016) were retrieved from NCBI SRA database. The short reads from the individual tissues were separately mapped to the genome to construct sequence alignments in HISAT2 (Kim et al. 2015). Subsequently, the reads were assembled and quantified using StringTie in terms of normalized transcripts per million (TPM) counts (Kovaka et al. 2019). Thereafter, the *AsCYPs* abundance in each tissue were measured and compared as (log TPM+1).

### 2.5. Homology Modelling and Substrate Identification of Sesquiterpene Oxidases

The protein sequences of *sesquiterpene oxidases* (expressed specifically in wounded wood tissue) were modelled based on the closely matched experimentally determined 3D structures in Protein Data Bank (PDB) using SWISS-MODEL server (Waterhouse et al. 2018). The substrate binding active site pocket was elucidated and defined based on the template structure using BIOVIA Discovery Studio software. The 3D structures of sesquiterpenes were retrieved from PubChem and Zinc20 database. Initially, the proteins were prepared by adding polar hydrogen and Kollman charges and energy of the substrate molecules were minimized, and grid box was generated surrounding the active site using AutoDock Tools. Subsequently, AutoDock Vina was used to perform molecular docking (Trott et al. 2010), PyMol to visualize the complex.

## 3. Results

### 3.1. Identification of Full-Length AsCYPs in *A. sinensis*

The *in-silico* analyses using the genome of *A. sinensis* enabled us to identify a total of 179 *AsCYP* genes possessing conserved P450 domains (**Table S1 and S2**). The gene names and their classification into families and subfamilies were designated based on the closest homology match of the experimentally verified CYPs of other plants from the Uniprot database. The percentage of protein sequence similarity with their homologs ranged from 45.48 to 94.86. The AsCYPs were classified into 8 clans and further subdivided into 42 families. Out of 42 families, 17 families (71, 73, 75, 76, 77, 78, 81, 82, 83, 84, 89, 92, 93, 98, 701, 703, and 736) belonged to A-type and 25 families (51 72, 74, 85, 86, 87, 88, 90, 94, 96, 97, 704, 707, 709, 710, 714, 715, 716, 720, 722, 721, 724, 734, 735, 749) belonged to non-A type. The CYP71 family had the most members, totaling 43. It was followed by the CYP81 family with 15 members and the CYP82 family with 14 members. However, four families, namely CYP715, CYP51, CYP74, and CYP720, consisted of only a single member each. The protein sequences within these families varied in length, with the number of amino acids ranging from 254 (AsCYP72A7) to 836 (AsCYP78A1). Furthermore, the molecular weights of the proteins spanned from 28.88 kDa (AsCYP72A7) to 92 kDa (AsCYP78A1). The isoelectric points (pI) also showed varied, ranging from 5.17 (AsCYP71BE10) to 9.9 (AsCYP72A5) (**Table S1**).

### 3.2. Phylogenetic Relations of AsCYPs in *A. sinensis*

To scrutinize the phylogenetic relations among the AsCYP superfamily members and with *A. thaliana* CYPs, a neighbor-joining (NJ) tree was drawn utilizing the alignment of the sequences of both the plant species. The tree with supported bootstrap percentage values revealed that AaCYPs were distributed and clustered into eight clans viz. 51, 71, 72, 74, 85, 86, 97, and 710 (**Fig. 1**). No AsCYP clustered with clan 711 member of *A. thaliana* which is consistent with the blast results, since no member belonging to the clan was detected. Within each clan, the members were sub-divided into families, where clan 71 possessed highest i.e., 17 families, followed by clan 85 (nine families), clan 72 (eight families), and clan 86 (four families) (**Table S3**), and they are regarded as multiple-family clans. The other four clans viz. clan 51, clan 74, clan 97, and clan 710 with single family each were designated as single-family clan. The clan 51, clan 74, and clan 710 had only one member each, while clan 71 had the highest (113), followed by clan 85 (27), clan 72 (17), clan 86 (16), and clan 97 (3). Within A-type clan, the family CYP71 had the highest members (43), followed by CYP81 (15), and CYP82 (14), while in non-A type clan, most members were from the families CYP72 (7), CYP716 (7) and CYP704 (5). Interestingly, in the tree, all the 179 AsCYPs were assembled into four major clusters. The clans viz., 51 and 71 stands out as two separate clusters, while clan 74, 85 and 710 clumped into single cluster. Similarly, clan 72, 86 and 97 grouped to form a different cluster. To be noted, the clan 71 was further sub-clustered into many groups consisting the members of similar families.

**Fig 1.**
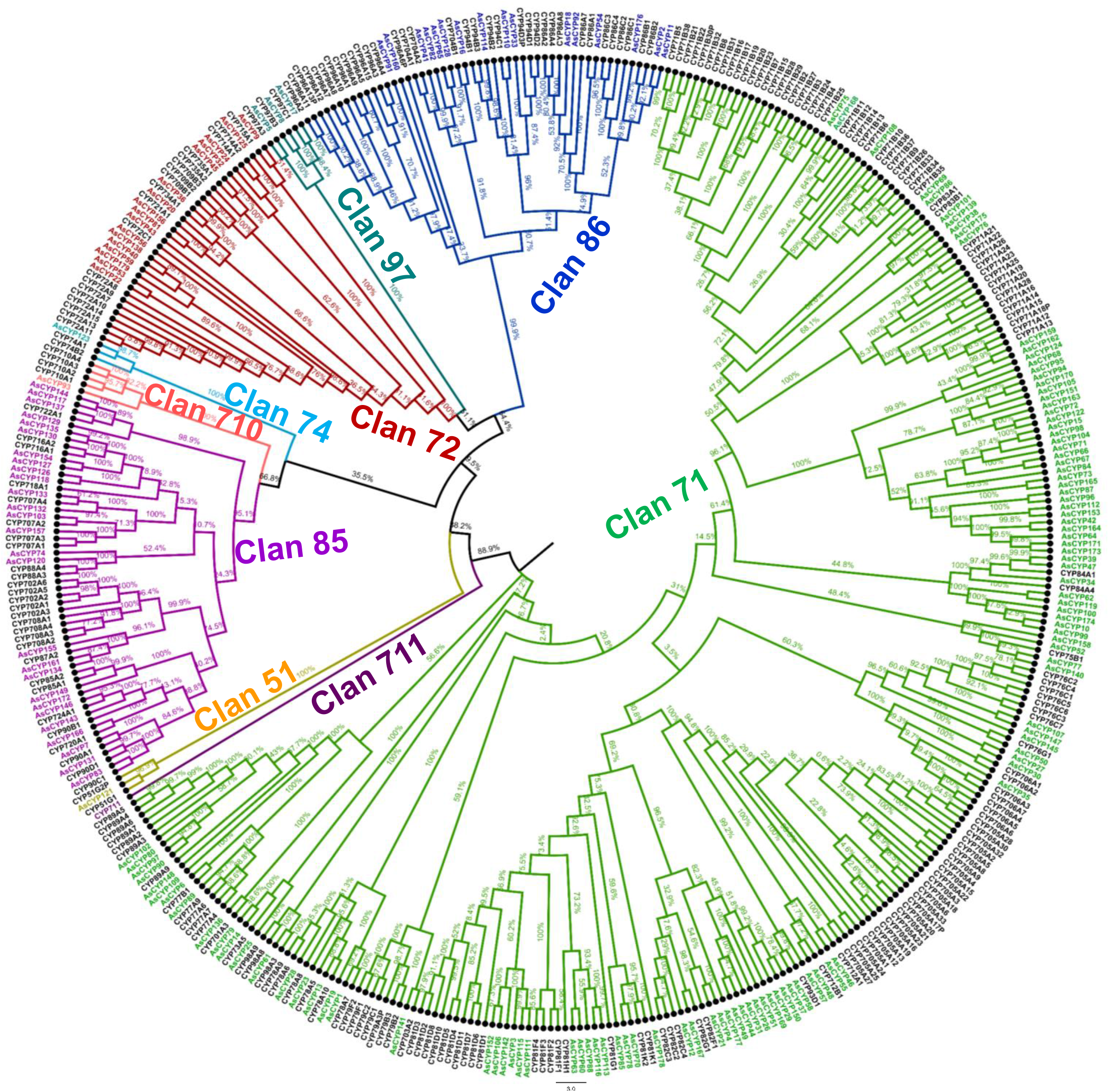
Phylogenetic positions of 179 *Aquilaria sinensis* CYPs (indicated in different color) with *Arabidopsis thaliana* CYPs (indicated in black) constructed using neighbor joining method in MEGA 7.0 using 1000 bootstrap. The AsCYPs classified into 8 clans were presented in different colors. The bootstrap percentage indicated reliability of the phylogenetic positions of the members.

To further elucidate the divergence and evolution of AsCYPs, a time tree was generated between *Arabidopsis thaliana*, *Oryza sativa*, *Vitis vinifera*, *Glycine max*, *Linum usitatissimum*, *Populus trichocarpa*, *Morus notabilis*, *Theobroma cacao*, *Solanum lycopersicum*, and *Gossypium raimondii*, and distribution of members into the clans was examined (**Fig. 2**). It was observed that emergence of the AaCYP superfamily in *A. sinensis* started after its divergence from *A. thaliana* around 110 million years ago (Mya). The species i.e., *A. thaliana*, *G. raimondii*, *T. cacao* and *A. sinensis* grouped into single clade where no clan 727 members were present but one each of clan 711 (except *A. sinensis*) were present. Additionally, *O. sativa* being monocot, appeared as single clade, and a greater number of clan 51 and 711 members were present compared to dicots. Comparatively, the count of clan 74 members was very less in *A. thaliana* and *A. sinensis* than other plant species. Interestingly, in all the species, including *A. sinensis*, clan 711 and 727 members were missing (**Fig. 2**). Additionally, in all the plants, the total gene count in the single-family clans was lower compared to the multiple-family clans. The cause of this disparity may be that the later have likely undergone expansion, facilitating the emergence of multiple subfamilies with specialized function within them.

**Fig 2.**
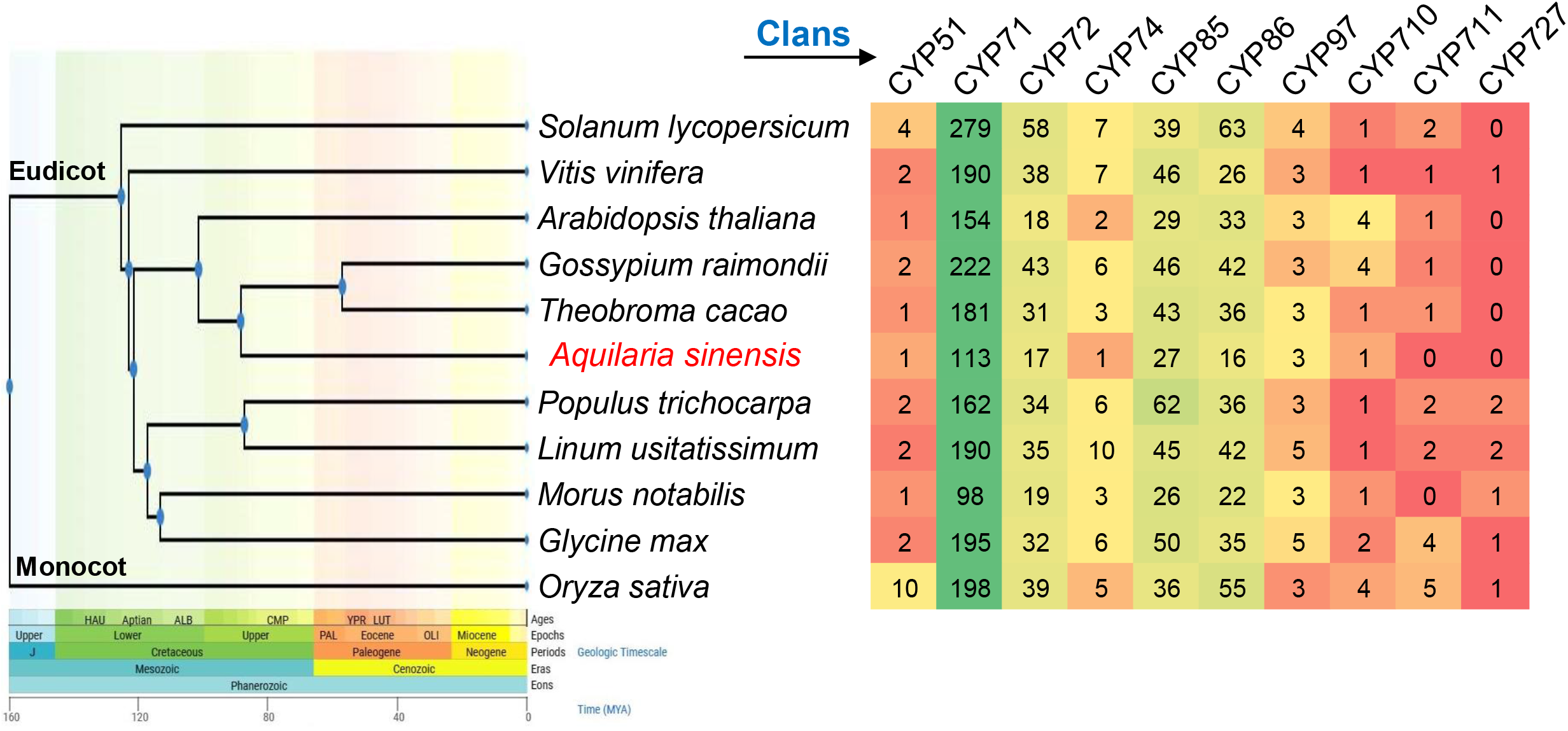
The distribution of number of CYPs of each clan in the genome of 11 plant species. The time tree indicates the diversification of the plant species with the time period in million years ago (Mya) and geological time scale. *Aquilaria sinensis* and *Arabidopsis thaliana* diversified from their common ancestor approximately 110 Mya.

### 3.3. Chromosomal Location and Clan Distribution

Physical loci of the 179 *AsCYP* genes were mapped to the eight chromosome (Chr) of *A. sinensis* and were found to be non-uniformly distributed (**Fig. 3**). Comparatively higher densities of *AsCYPs* were located in the upper and middle arm of the chromosomes. The number of genes varied in each Chr. For instance, Chr01 had the highest number of *AsCYPs*, followed by Chr06 (27), Chr02 (26), and Chr05 (23). Interestingly, Chr03 and Chr07, in spite of difference in length, had the same number of genes (21). However, Chr08 being the smallest in length had only eight *AsCYPs*. Therefore, Pearson correlation coefficient value was calculated to verify if any correlation between the length of chromosomes and number of genes exists (**Fig. 4**). A high positive correlation (r = 0.80, p value = 0.016) suggested that considering the number of *AsCYPs*, their distribution of in the Chr of *A. sinensis* was not random. However, the clan members, particularly those contained multiple families were randomly distributed (**Fig. 3**). For e.g., members of two clan (71 and 85) were located in all the 8 chromosomes, while clan 72 members were absent in Chr01 and Chr02. Similarly, clan 86 was absent in Chr03 and Chr04. Moreover, the single-family clans viz. 51, 74 and 710 with only single member each were restricted to Chr01, Chr01, and Chr03, respectively. Alternately, the clan 97 members, 3 in total, were dispersed in Chr06 and Chr07. Overall, distribution pattern of the *AsCYPs* indicated that their expansion possibly occurred through whole genome duplication events.

**Fig 3.**
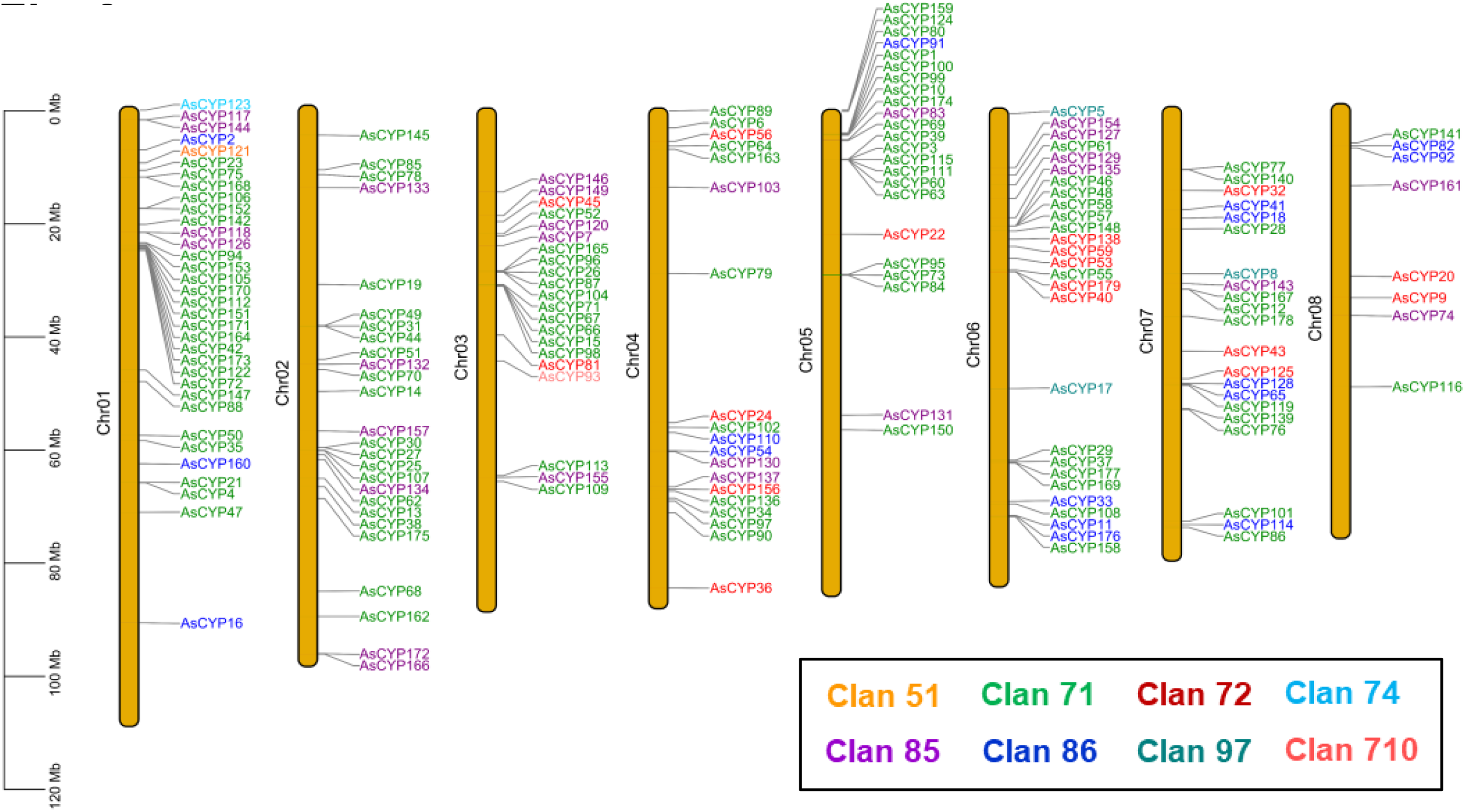
Chromosomal location of 179 *AsCYPs* in *A. sinensis*. The scale on the left side represents 120 Mb chromosomal distance, and the grey bar corresponding to each *AsCYP* represent its physical location. The clan members are presented in different colors viz. yellow (clan 51), green (clan 71), red (clan 72), cyan (clan 74), violet (clan 85), blue (clan 86), dark-blue (clan 97), and pink (clan 710).

**Fig 4.**
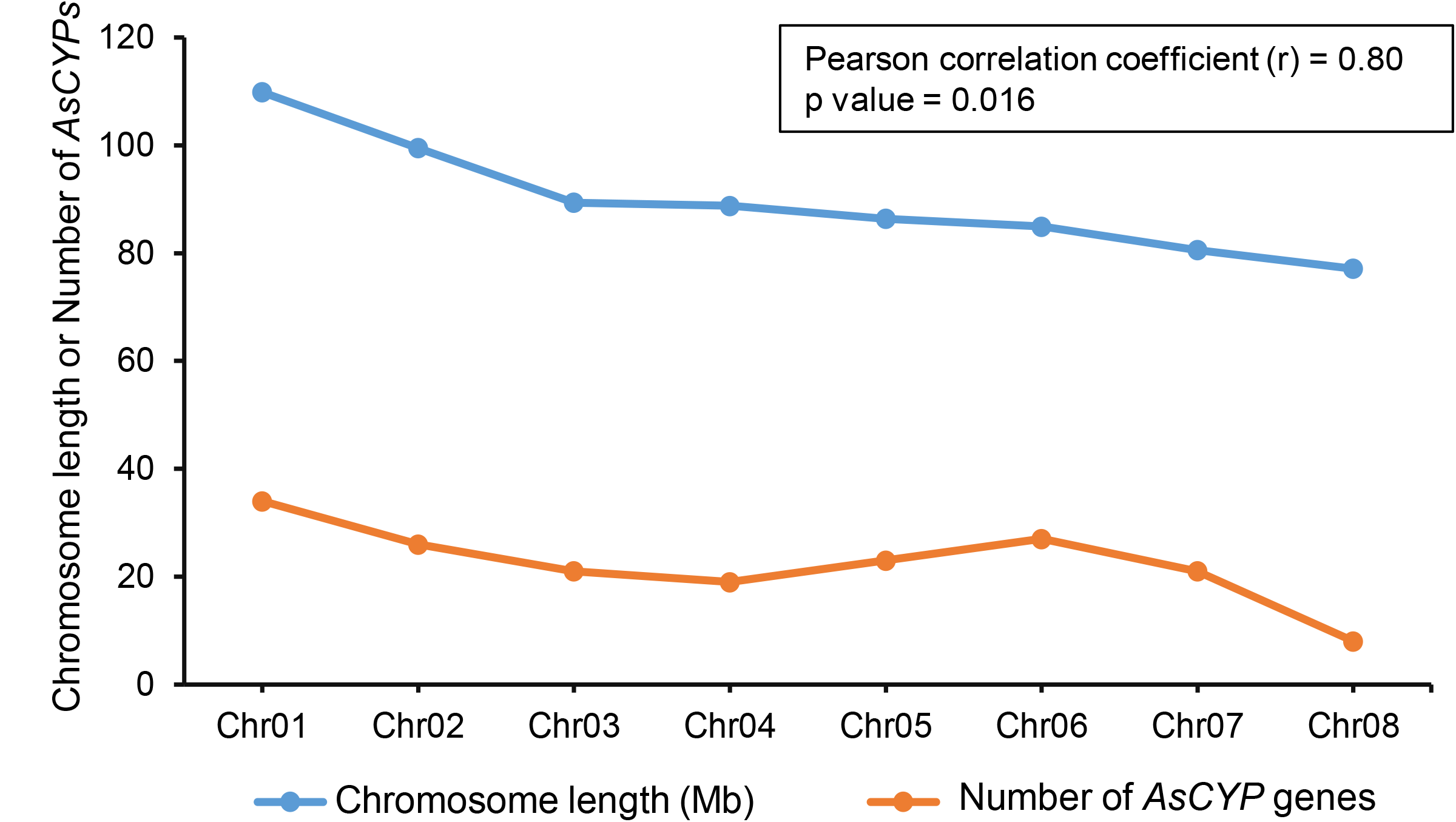
Chromosome distribution of *AsCYPs* in *Aquilaria sinensis*. The positive correlation of number of AsCYP genes and the length of the 8 chromosomes into which they are located, verified by Pearson correlation coefficient value.

### 3.4. Duplication and Selection Pressure of *AsCYP* genes

To better understand the evolution and expansion of the AsCYPs in the genome, duplication analysis was conducted within the *A. sinensis* genome. This study also enabled us to identify the paralogous gene pairs. Out of the 179 *AsCYPs*, 51 (28.4%) were found to be duplicated due to segmental duplication (SD), while only 31 (17.31%) were duplicated through tandem duplication (TD) events. This resulted in generation of 44 SD gene pairs and 23 TD gene pairs (**Table S4**). Interestingly, 14 genes were associated with both SD and TD events. The tandem duplicated genes were restricted to 5 chromosomes (Chr01, Chr02, Chr03, Chr05, and Chr06), while the segmentally expanded genes were dispersed throughout all 8 chromosomes **(Fig. 5)**. The majority of duplicated members (50) belonged to clan 71, followed by clan 85 (7), clan 72 (6), and least to the clan 86 (4). It is worth noting that among all the SD genes, *AsCYP71BE5* and *AsCYP72A6*, as well as among the TD genes, *AsCYP71AU3* and *AsCYP71AU1*, exhibited the most duplicated pairs but they expanded into same families, indicating their significant functional role in *A. sinensis*.

**Fig 5.**
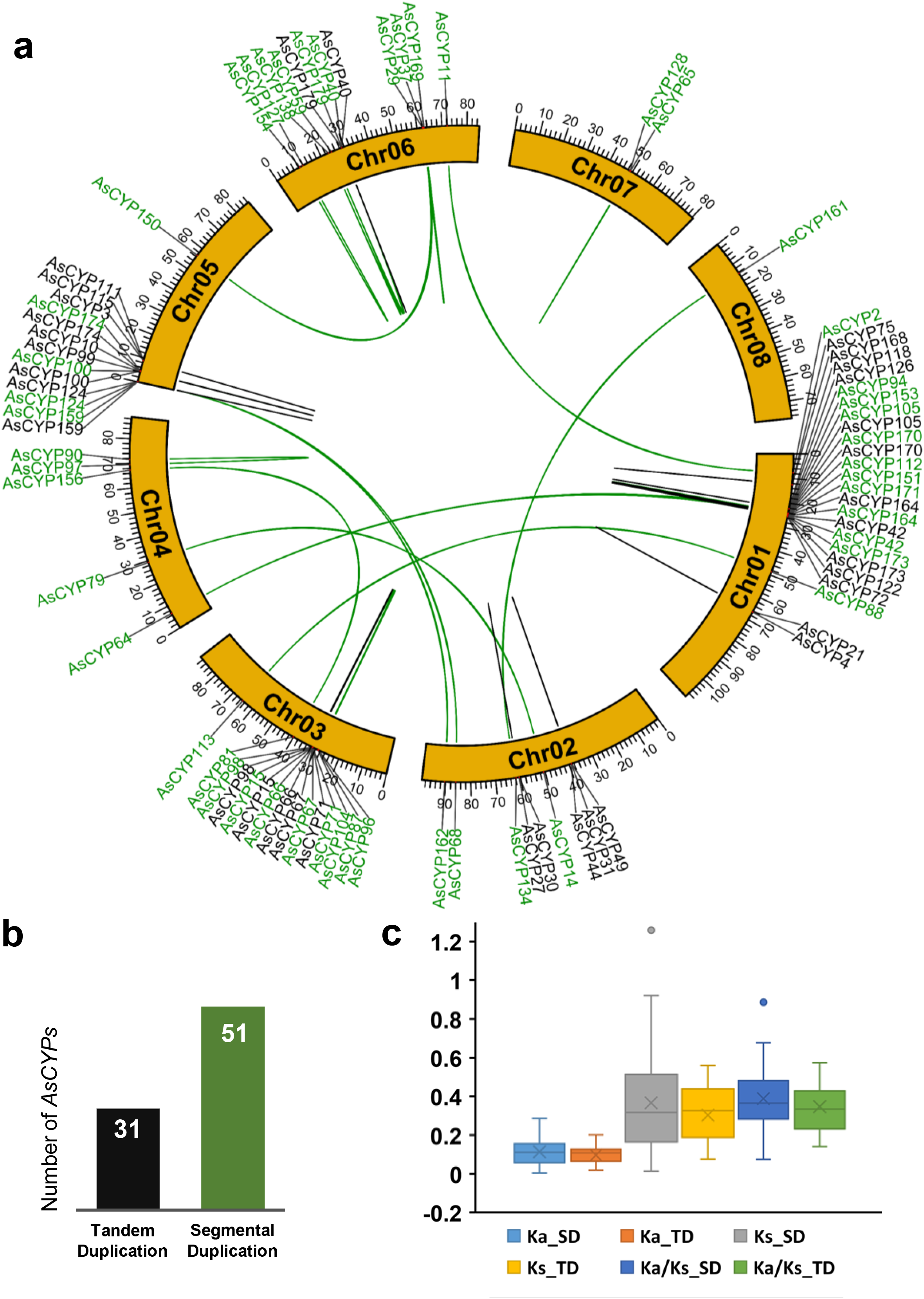
The expansion and evolution of AsCYPs in *A. sinensis*. **(a)** Distribution of the duplicated gene in the 8 chromosomes. The black and green lines represent the tandem and segmental duplicated gene pairs. **(b)** Bar graph representing the total number of tandem and segmental duplicated genes in the genome. **(c)** Box plot representing the Ka, Ks, and Ka/Ks values of AsCYPs undergoing different duplication modes where SD stands for segmental duplication and TD stands for tandem duplication

Furthermore, collinearity and syntenic analysis were conducted to identify evolutionary related orthologous genes between *A. sinensis* and *A. thaliana*. In total, 21 gene were identified which were distributed across all chromosomes except Chr01. Among them, Chr04 and Chr05 exhibited the highest number of orthologous pairs with 5 pairs each, followed by Chr02, Chr03, and Chr07 with 3 pairs each. Chr06 and Chr08 had the lowest number of orthologous pairs, with only 1 pair each (**Fig. 6**). Contrastingly in *Arabidopsis*, the orthologous members were scattered throughout all 5 chromosomes. Most of the orthologous *AsCYPs* were from the clan 71 (8), followed by clan 85 (7), clan 86 (6), and clan 72 (1). Similar to the paralogous pairs, the orthologous members (except *AsCYP85A1*) displayed divergence and expansion into members of similar families indicating evolutionary related conserved functional role (**Table S5**). To further examine the nature and time of evolution, Ka, Ks, and Ka/Ks ratios of the duplicated *AsCYPs* were evaluated (**Table S6**). In some pairs, Ks values were not obtained due to high sequence divergence, possibly attributed to codon biases or alignment issues, and these pairs were excluded from the analysis. The average Ks values for SD and TD *AsCYPs* was 0.36 and 0.30, respectively. Whereas the average Ka values for SD was 0.01 and TD was 0.09. The study found that the mean divergence time for SD genes was approximately 26.15 Mya, while for TD genes, it was around 21.64 Mya. The mean Ka/Ks ratios which indicated the selection pressure on the genes, were found to be 0.38 for SD genes and 0.34 for TD genes. The Ka/Ks ratios of the orthologous pairings were within narrow range i.e., from 0.046 (*AsCYP77B1*/*AT1G11600.1*) to 0.369 (*AsCYP85A1*/*AT5G14400.1*), whereas the paralogous pairs showed a wide range of values i.e., from 0.073 (*AsCYP73A1*/*AsCYP73A2*) to 0.893 (*AsCYP704C2*/*AsCYP704C4*). To note, the Ka/Ks ratios of all the duplicated *AsCYPs* gene pairs irrespective of their duplication mode were <1.

**Fig 6.**
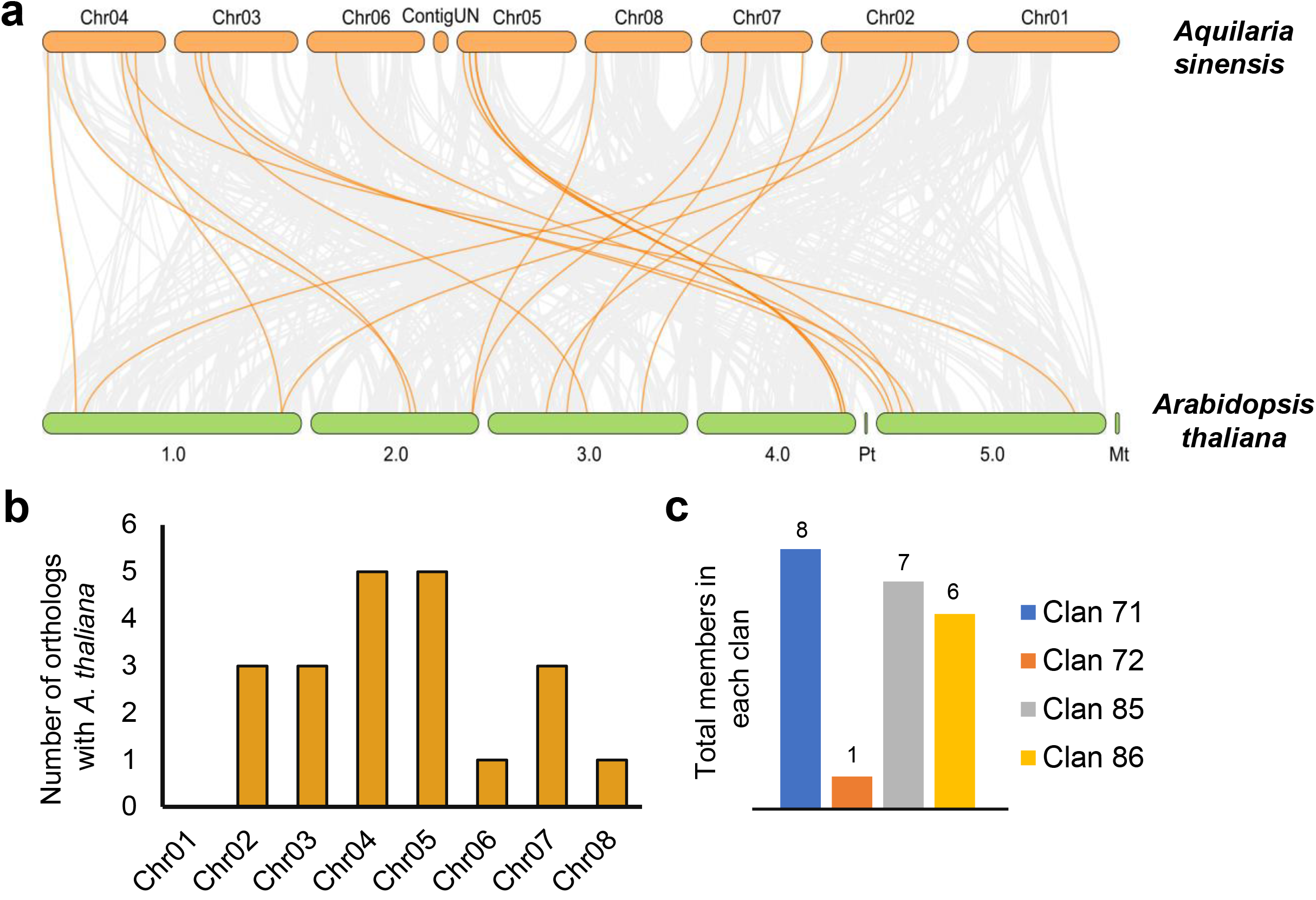
Collinearity and synteny analysis between *A. sinensis* and *A. thaliana*. **(a)** Syntenic blocks representing the evolutionary related coding regions between *A. sinensis* and *A. thaliana*. The yellow lines represent syntenic relations in terms of the genomic positions and dispersion of evolutionary related CYPs of *A. sinensis* and *A. thaliana*. **(b)** Bar plot representing the distribution of total number of orthologous genes in the chromosomes of *A. sinensis*. **(c)** Bar plot representing the distribution of orthologous genes into CYP clans.

### 3.5. Functional Positions of AsCYP Superfamily in *A. sinensis*

CYPs are directly involved various biological and molecular processes, including the secondary metabolites biosynthesis which are essential for physiology and growth of plants. To understand the functional role of AsCYP proteins, GO and InterPro terms were assigned, and were explored in KEGG and UniProt database for functional annotation. Homology-based functional positions were obtained and summarized (**Table S7**). The evidence gathered corroborate their involvement in the synthesis of diverse secondary metabolites including terpenoids, flavonoids, isoflavonoids, phenylpropanoids, alkaloids, carotenoids, zeatin, lipids (wax, suberin, cutin, steroids, alpha-linolenic acid), and hormones (jasmonate, gibberellin, brassinosteroid, and abscisic acid). These results further suggest their participation in the production of vitamins like quinones, as well as other natural compounds including sapogenin, glucosinolate, methoxypsoralens, furonapththodioxoles, phenylurea, (seco)iridoids, pterocarpan, biphenyl, and cyanogenic glycosides. Notably, a few AsCYPs were found to be involved in functions such as volatile emission and wound response. Furthermore, it was noted that the count of AsCYPs involved in sesquiterpenoids biosynthesis was highest (30), followed by diterpenoids (15) and isoflavonoid (12) biosynthesis. A few were associated with processes such as hormone signaling (10), fatty acid metabolism (8), brassinosteroid (8), and cyanogenic glycoside (8) biosynthesis (**Fig. 7**). Overall, the functional positions of the AsCYPs and the significant number of genes specifically associated with these pathways indicate their collective role and involvement in hormone-mediated regulation of secondary metabolites, possibly during wounding and stress responses in *A. sinensis*.

**Fig 7.**
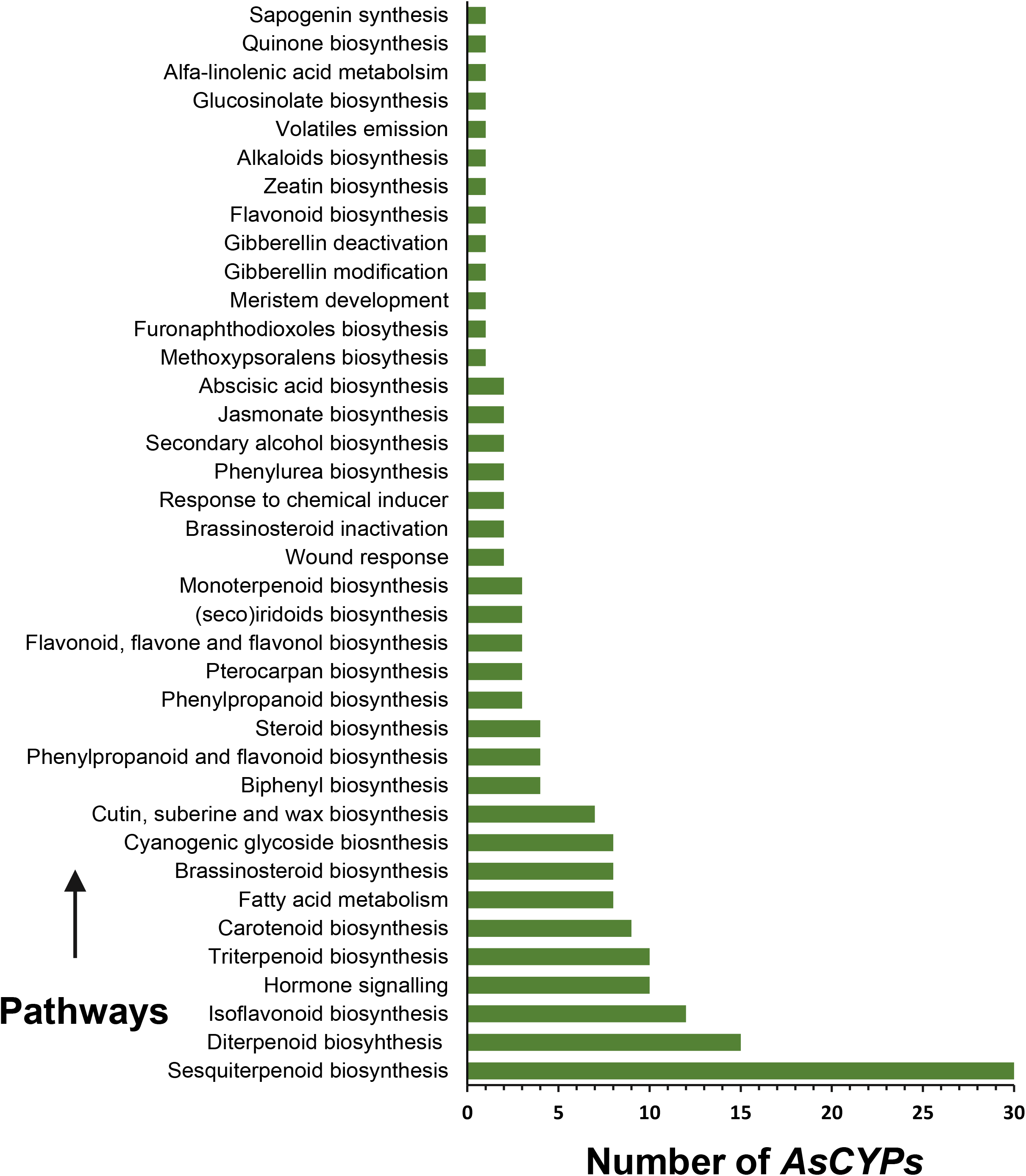
Functional role of the AsCYPs analysed from Uniprot and KEGG annotation. The X-axis represents the number of AsCYPs and Y-axis represents the pathways they are involved.

### 3.6. Expression Profile of *AsCYPs* in Multiple Tissues of *A. sinensis*

To measure the transcript levels of *AsCYPs* in 11 different types of tissues, including wounded wood (Ordinary and Chi-Nan germplasm) and stem, leaves, flowers, buds, aril, seeds, and salt-treated callus tissue, transcriptomic reads were extracted. The average counts were then converted into logarithmic TPM expression values, and hierarchical clustering was performed (**Fig. 8 and Table S8**). Based on their expression patterns, the *AsCYPs* were categorized into 8 clusters. In group 1, 21 *AsCYPs* exhibited relatively higher expression levels in at least 8 tissues. Among them, two *AsCYPs*, namely *AsCYP73A1* and *AsCYP73A2*, which encoded trans-cinnamate 4-monooxygenase, consistently showed higher expression across all tissues. Additionally, *AsCYP71D12* and *AsCYP71D15* (encoding alpha-guaiene 2-oxidase) exhibited elevated transcript levels in both types of wounded and healthy wood, stem, callus, seed, and flower. Group 2 consisted of *AsCYPs* that showed higher expression in callus, wounded stem, wood, aril, seed, flower, and bud when compared to other tissues. The members in this group encoded proteins such as alpha-guaiene 2-oxidase (AsCYP71BE2, AsCYP71BE5, AsCYP71D4, AsCYP71D9), sterol 22-desaturase (AsCYP710A1), isoflavone 2’-hydroxylase (AsCYP81Q2), and those involved in jasmonate and ethylene signaling (AsCYP82A10). In group 3, out of the 39 *AsCYPs*, 24 genes showed moderate expressions in all the tissues. However, two genes, *AsCYP71A2* and *AsCYP701A1*, encoding dammarenediol 12-hydroxylase and ent-kaurene oxidase, respectively, exhibited higher expression in both wounded and control wood of Ordinary and Chi-Nan *A. sinensis*. Furthermore, in the same group, *AsCYP71D6* (encoding alpha-guaiene 2-oxidase) displayed higher expression specifically in wounded wood, indicating its tissue-specific nature. Transcripts of the *AsCYPs* in group 4 showed relatively elevated levels in wounded stem, wounded Chi-Nan wood, leaves, flowers, and buds compared to other tissues. This group included genes encoding geraniol 8-hydroxylase (*AsCYP76B2*), fatty acid omega hydroxylase (*AsCYP86A1*), flavonoid 3’,5’-hydroxylase (*AsCYP75A2*), alpha-guaiene 2-oxidase (*AsCYP71D10*), and isoflavone 2’-hydroxylase (*AsCYP81Q5*). Interestingly, group 5 consisted of four tissue-specific genes that were exclusively expressed in seeds. Among them, three genes (*AsCYP71D8*, *AsCYP71BE4*, and *AsCYP71BE6*) encoded alpha-guaiene 2-oxidase, while one gene (*AsCYP716B2*) encoded abietadienal oxidase. Group 6 and 7 genes displayed moderate and low expression levels, respectively, across all tissues compared to the other groups. Conversely, the genes in group 8 exhibited relatively higher expression in wounded stems and callus tissue. Overall, as per observation, there was variation in the pattern of expression even between the same family members, indicating functional divergence among themselves. Some genes showed tissue-specific expression, while a few exhibited widespread expressions across all tissues.

**Fig 8.**
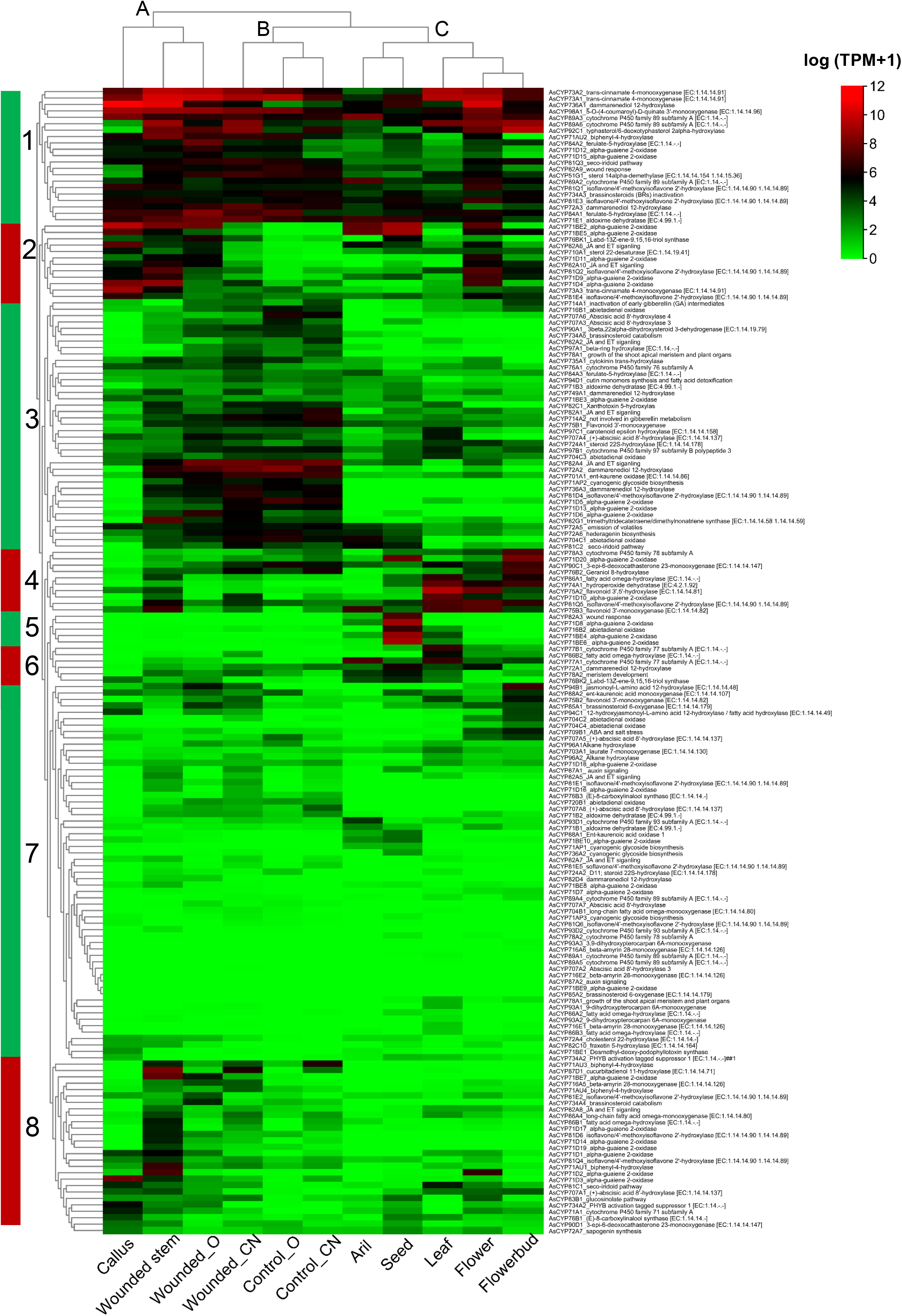
Expression profiles of *AsCYPs* in different tissues of *A. sinensis*. Mean expression values of the genes in each tissue were considered for data normalization which were converted to log base of 2. Green, black and red bar represents low, moderate and high expressions, respectively. Horizontal clustering is grouped into 8 major clusters based on the individual expression pattern of the genes, and vertical clustering is grouped into 3 cluster based on transcript abundance in each tissue.

Furthermore, it is noteworthy that the highest number of *AsCYPs* with higher transcript abundance was observed in wounded Chi-Nan wood, followed by wounded Ordinary wood and wounded stems of *A. sinensis*. The clustering of the tissues (columns) resulted in three major clusters (A, B, and C) (**Fig. 8**). Within cluster A, wounded stem and Ordinary wood were grouped together, while the callus tissue stood out. In cluster B, both control wood samples grouped together, with Chi-Nan wood standing out. Similarly, in cluster C, as expected, the aril and seed formed one group, while the flower, bud, and leaf formed another. The tissues that fell into separate clusters indicated differences in the level of expression of the *AsCYPs*, while tissues in the same sub-clusters exhibited similar expression patterns.

### 3.7. Substrate Prediction of a Sesquiterpene Oxidase Encoded by *AsCYP71D6*

It was noted that the gene *AsCYP71D6* which encoded a sesquiterpene oxidase (SO), exhibited specific elevation in wounded wood tissue. Considering the substrate selectivity and versatility of CYPs, as well as the presence of numerous oxygenated sesquiterpene derivatives profile reported in agarwood, we decided to expand our study and explore the potential substrates of the enzyme. This was accomplished by performing 3D modeling to identify the active pocket of AsCYP71D6, followed by screening and validating 395 sesquiterpenes for their binding affinity through molecular docking and interaction analysis. The 3D structure of AsCYP71D6 which was modeled based on the template structure (PDB:5YLW), comprised of 22 α-helices, 4 β-strands, turns, and loops, as depicted in **Fig 9**. The Root Mean Square Deviation (RMSD) value of 0.206 Å, representing the alignment score between the constructed model and the template, indicated that the model had an acceptable quality. The sesquiterpene molecules docked into the active pocket of AsCYP71D6, allowed us to identify 11 potential substrates, namely isovalencenol, kusunol, eudesmol, suberosol D, germacrene D, kunzeaol, b-betulenol, eudesm-7(11)-en-4-ol, alloeremophilone, eremophiladien-12-oic acid, and preisocalamendiol. The substrates were considered according to their binding energy and interaction with the active site residues. The docking analysis revealed a binding affinity range of -7.7 to -7.3 kcal/mol, indicating stable interactions within the enzyme-substrate complex. Notably, conventional hydrogen bonds and alkyl bonds were observed, signifying favorable binding interactions. (**Table 1**). Moreover, within the active site pocket, the residues GLU:302, THR:303, and GLN:295 were engaged in hydrogen bonding interactions, while ALA:118, PHE:119, PHE:238, ALA:480, LEU:368, CYS:438, LEU:364, and PRO:363 participated in alkyl bonding interactions. LEU:298 played a role in both types of bonding, while the other residues within the pocket likely facilitated hydrophobic interactions, further enhancing the stability of the complex.

**Fig 9.**
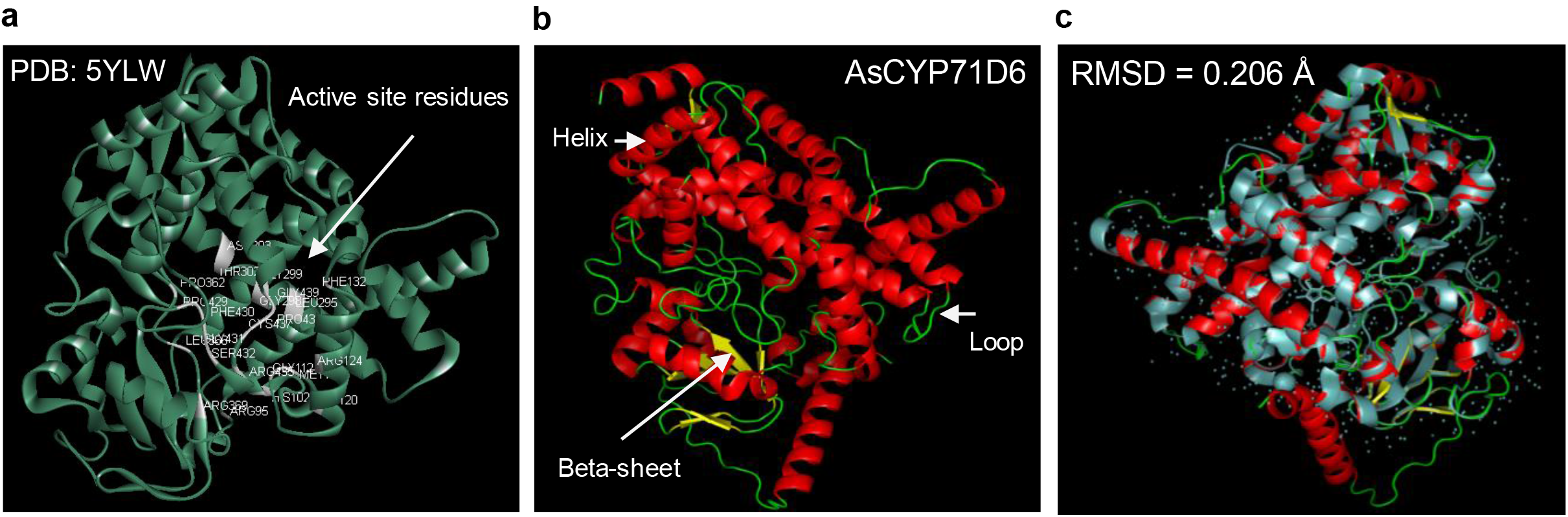
3D modelling of the sesquiterpene oxidase (AsCYP71D6) of A. sinensis. **(a)** Template 3D structure (PDB: 5YLW) based on it the structure of AsCYP71D6 was constructed. The active site residues representing in white color is defined by analyzing the substrate interaction with the pocket. **(b)** The model of AsCYP71D6 consisting of helices (red), beta-sheets (yellow) and loops (green). **(c)** 3D alignment of the template and model with RMSD value of 0.206 Å indicate good quality of the model.

**Table 1.**
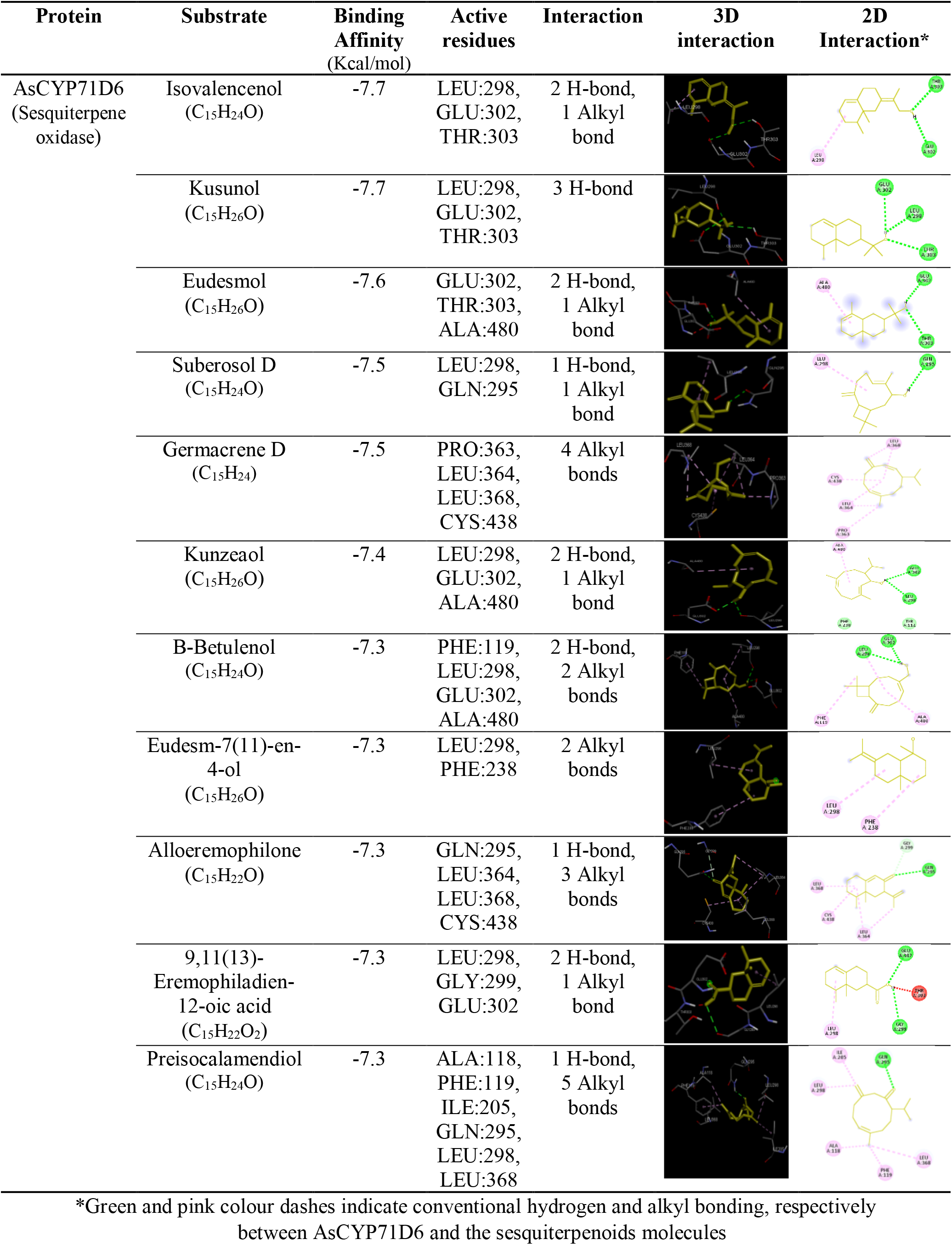
List of Possible substrates of AsCYP71D6 identified through molecular docking and interaction analysis.

| Protein | Substrate | Binding Affinity (Kcal/mol) | Active residues | Interaction | 3D interaction | 2D Interaction* |
| --- | --- | --- | --- | --- | --- | --- |
| AsCYP71D6<br>(Sesquiterpene oxidase) | Isovalencenol<br>(C<sub>15</sub>H<sub>24</sub>O) | -7.7 | LEU:298,<br>GLU:302,<br>THR:303 | 2 H-bond,<br>1 Alkyl bond |  |  |
|  | Kusunol<br>(C<sub>15</sub>H<sub>26</sub>O) | -7.7 | LEU:298,<br>GLU:302,<br>THR:303 | 3 H-bond |  |  |
|  | Eudesmol<br>(C<sub>15</sub>H<sub>26</sub>O) | -7.6 | GLU:302,<br>THR:303,<br>ALA:480 | 2 H-bond,<br>1 Alkyl bond |  |  |
|  | Suberosol D<br>(C<sub>15</sub>H<sub>24</sub>O) | -7.5 | LEU:298,<br>GLN:295 | 1 H-bond,<br>1 Alkyl bond |  |  |
|  | Germacrene D<br>(C<sub>15</sub>H<sub>24</sub>) | -7.5 | PRO:363,<br>LEU:364,<br>LEU:368,<br>CYS:438 | 4 Alkyl bonds |  |  |
|  | Kunzeaol<br>(C<sub>15</sub>H<sub>26</sub>O) | -7.4 | LEU:298,<br>GLU:302,<br>ALA:480 | 2 H-bond,<br>1 Alkyl bond |  |  |
|  | B-Betulenol<br>(C<sub>15</sub>H<sub>24</sub>O) | -7.3 | PHE:119,<br>LEU:298,<br>GLU:302,<br>ALA:480 | 2 H-bond,<br>2 Alkyl bonds |  |  |
|  | Eudesm-7(11)-en-4-ol<br>(C<sub>15</sub>H<sub>26</sub>O) | -7.3 | LEU:298,<br>PHE:238 | 2 Alkyl bonds |  |  |
|  | Alloeremophilone<br>(C<sub>15</sub>H<sub>22</sub>O) | -7.3 | GLN:295,<br>LEU:364,<br>LEU:368,<br>CYS:438 | 1 H-bond,<br>3 Alkyl bonds |  |  |
|  | 9,11(13)-Eremophiladien-12-oic acid<br>(C<sub>15</sub>H<sub>22</sub>O<sub>2</sub>) | -7.3 | LEU:298,<br>GLY:299,<br>GLU:302 | 2 H-bond,<br>1 Alkyl bond |  |  |
|  | Preisocalamendiol<br>(C<sub>15</sub>H<sub>24</sub>O) | -7.3 | ALA:118,<br>PHE:119,<br>ILE:205,<br>GLN:295,<br>LEU:298,<br>LEU:368 | 1 H-bond,<br>5 Alkyl bonds |  |  |

\*Green and pink colour dashes indicate conventional hydrogen and alkyl bonding, respectively between AsCYP71D6 and the sesquiterpenoids molecules

## 4. Discussion

The Cytochrome P450s (CYPs) are a superfamily of heme-thiolate class of enzymes that facilitate a wide range of chemical reactions. These enzymes are found in all kingdoms of life and have especially experienced significant diversification within the plant kingdom. CYPs have been identified at the genome level in a number of plant species, including *Arabidopsis thaliana* (Xu et al. 2001), *Vitis vinifera* (Jiu et al. 2020), *Glycine max* (Guttikonda et al. 2010), *Ricinus communis* (Kumar et al. 2014), *Linum usitatissimum* (Babu et al. 2013), *Oryza sativa* (Wei at al. 2018), *Populus trichocarpa* (Nelson et al. 2008), *Morus notabilis* (Ma et al. 2014). In a previous study, we successfully identified, characterized, and functionally classified the superfamily members in *Aquilaria agallocha*, validating the functions of a few members associated with terpenoids and phenylpropanoids biosynthesis (Das et al. 2023). However, comprehensive analysis regarding the evolution and tissue-wide expression of the superfamily in *Aquilaria* is missing. In this study, our main objectives were to identify and gain insights into the evolutionary relations, including duplication and selection pressure behind the expansion of the AsCYPs. Additionally, to estimate the expression patterns of the members in the RNA-seq data of various tissues, including ordinary and Chi-Nan agarwood.

### 4.1. CYP Superfamily in *Aquilaria sinensis* and their Evolution

In the present study, 179 full-length CYPs were identified and classified into 8 clans and 42 families in *A. sinensis*. The presence of considerable number of CYPs in the genome of various organisms indicate their early diversification during evolution. For instance, even in simpler organisms such as lycophyte *Selaginella moellendorffii* (199), hornwort *Anthoceros agrestis* (144), moss *Physcomitrella patens* (71), liverwort *Marchantia polymorpha* (115), algae *Klebsormidium nitens* (29), as well as in angiosperm monocot like *Oryza sativa* (329 CYPs), and angiosperm eudicot like *Arabidopsis thaliana* (246), a significant number of CYPs were found (Nelson et al. 2009). The widespread distribution across different organisms suggest that CYPs have been subjected to diversification and expansion throughout evolutionary journey. In comparison to *A. sinensis*, *A. agallocha* exhibits a smaller repertoire with only 38 families, lacking six families specifically: CYP72, CYP92, CYP93, CYP709, CYP721, and CYP749 (Das et al. 2023). The absence of certain families in *A. agallocha* could be attributed to two possible reasons. Firstly, it could be due to errors in the genome assembly, which might require further refinement and improvement to accurately identify all the CYP families. Secondly, over time, few CYPs may have undergone evolutionary changes, causing their loss or divergence into other families. Additionally, the member of the families viz., CYP79, CYP705, CYP706, CYP708, CYP702, CYP718, CYP711, and CYP712 were missing in *A. sinensis* but present in *A. thaliana* (Nelson et al. 2009). Interestingly, three families (CYP92, CYP736, CYP749) were exclusively present in *A. sinensis* but not in *A. thaliana.* These members may have emerged to carry out specific functions in *A. sinensis.* Within dicots, there was notable differences in the number of *CYPs* identified. The number of genes in *A. sinensis* was lower compared to *Vitis vinifera* (457), *Eucalyptus grandis* (443), *Glycine max* (332), *Oryza sativa* (329), and *A. thaliana* (246) (Nelson and Werck-Reichhart 2011, Nelson et al. 2009). But higher than *Morus notabilis* (174) and *A. agallocha* (136). The presence or absence of specific family members among different plant species indicates variations in their phytochemical profiles and, consequently, in stress adaptation mechanisms (Xu et al. 2015). Moreover, it is interesting to note that the CYPomes (the complete set of CYP genes) in the genomes of different plant species exhibit variations in relation to genome size and protein coding genes (PCGs). *A. thaliana*, with a 133 Mb of genome, the CYPomes constitute approximately 0.93% of the PCGs. While in *A. sinensis* (726 Mb), *A. agallocha* (726 Mb) and *O. sativa* (321 Mb), having larger genomes than *A. thaliana*, CYPomes constitute approximately 0.92%, 0.61%, and 0.53% of the PCGs, respectively (Wang et al. 2021, Kawahara et al. 2013, Das et al. 2021). However, it is worth mentioning that *M. notabilis*, despite having a smaller genome size of 330 Mb compared to *Aquilaria* species, exhibits a similar proportion of CYPomes i.e., approximately 0.59% of the PCGs (He et al. 2013). These discrepancies in genome size, PCGs, and CYPomes among plant species highlight the unique evolutionary pathways or variation in duplication events taken by different plants in adapting to specific environmental challenges and optimizing their response to various stressors. It further suggests that the distinct sets of CYP family members in the plant species enables them to effectively cope with specific environmental conditions and ensure their survival and adaptation. In *A. sinensis*, existence of considerably higher number of *AsCYPs* associated with sesquiterpenoids pathway indicates the importance of these enzymes in producing sesquiterpenoids and probably reflects the plant’s ability to respond and adapt to wounding or injury (Xu et al. 2013). Nevertheless, despite the variation, we observed that 21 *AsCYPs* in *A. sinensis* shared an orthologous relationship with *CYPs* found in *A. thaliana*. This finding suggests that these members might have been conserved during speciation and could potentially share close functional relations.

The distribution and clustering of AsCYPs into 8 clans within the evolutionary tree provided insights into their evolutionary history. Certain clans (51, 74, 86, and 97), remained conserved across plants and shared orthologous functional relationship (Hansen et al. 2021). Clan 51, which existed as a separate cluster, likely originated from an ancestor associated with sterol metabolism and was involved in steroid biosynthesis in *A. sinensis* (Kim et al. 2005). Clan 71, on the other hand, had evolved and expanded into the highest number of families, indicating significant functional divergence associated with the biosynthesis of metabolites such as sesquiterpenoids, flavonoids, and phenylpropanoids (Hamberger et al. 2013, Yonekura-Sakakibara et al. 2019, Hansen et al. 2021). The appearance of clans 74, 85, and 710 in the similar cluster suggested that their members were generated from a single ancestor. In addition, clan 74 was involved in jasmonic acid and oxylipin biosynthesis, while clan 85 in brassinosteroid (BRs) biosynthesis, and clan 710 in membrane sterol biosynthesis (Sivasankar et al. 2000, Castle et al. 2005, Morikawa et al. 2006). Similarly, clans 72, 86, and 97 had also evolved from common ancestors, where clan 72 members were involved in the metabolism of isoprenoid compounds, including diterpenes and triterpenes, as well as alkaloid metabolism. Clan 86 and 97 was associated with biosynthesis of fatty acid and carotenes, respectively (Hansen et al. 2021, Han et al. 2010, Kim et al. 2009). Overall, the evolutionary tree extended firm indication that AsCYPs had undergone duplication and expansion from common ancestors, giving rise to different clan and family members with functional divergence. This diversification and specialization of CYPs is likely to play crucial role in the synthesis of a diverse range of secondary metabolites in *A. sinensis*.

### 4.2. The AsCYP Superfamily Expanded Through Segmental Events under Negative Selection Pressure

Gene duplication has remained as a primary tool exploited by plant genomes to acquire novel functions, leading to divergence and, ultimately, speciation (Cannon et al. 2004). We observed an uneven distribution of the 179 AaCYPs belonging to 8 clans and 42 families in the 8 chromosomes. Gene duplication study revealed that most of the *AsCYP* genes evolved as a result of segmental duplication (SD) events, implying a non-rapid or slow evolutionary process, as SD was the primary force for their expansion. This finding was consistent with *O. sativa*, *G. max*, and *A. thaliana*, where most of the CYPs arose from SD events (Wei at al. 2018, Yu et al. 2017, Guttikonda et al., 2010). However, it contrasted with *V. vinifera*, where CYPs mainly arose due to tandem duplication (TD) events (Jiu et al. 2020). The fluctuation in values of Ka, Ks, and Ka/Ks of the duplication modes suggest variation in the evolution rates of the genes (Que et al. 2018). The higher Ks values of *AsCYP* SD genes suggested that they likely underwent evolution earlier and have more accumulated changes over time. In addition, the ratio (Ka/Ks) of both SD and TD genes being < 1 indicated purifying or negative selection pressure acting on the *AsCYP* genes. Moreover, the high values of Ks were reflected in the divergence time as well. After the divergence of *A. thaliana* and *A. sinensis* approximately 110 Mya, the first SD event occurred between *AsCYP87A1* and *AsCYP87A2* around 90.47 Mya, while the most recent SD event occurred between *AsCYP734A3* and *AsCYP734A5* approximately 0.95 Mya. The series of duplications occurred between this time period had likely facilitated the expansion and diversification of the CYPs in genome of *A. sinensis*.

### 4.3. Variation in the expression level of *AsCYPs* among different tissues may linked to functional divergence

Plant CYPs takes part in numerous biochemical reactions and their diverse physiological roles has been previously illustrated (Werck-Reichhart and Feyereisen 2000). These versatile enzymes play crucial roles in the biosynthesis of various secondary metabolites, including terpenoids, flavonoids, phenylpropanoids, alkaloids, carotenoids, and hormones like jasmonate, gibberellin, brassinosteroid, and abscisic acid (Hansen et al. 2021). In this study, we have also observed similar findings regarding the functional diversity of AsCYP members, as illustrated through KEGG mapping. The results provide further evidence of the essential roles played by CYPs and diverse biochemical processes of *A. sinensis*. Moreover, in plants including Arabidopsis, rice, common grape and soyabean, expression pattern of candidate CYPs in various tissues including vegetative, stress induced and developmental stages have been studied to elucidate their tissue specific functional roles (Xu et al. 2001, Wei at al. 2018, Jiu et al. 2020, Guttikonda et al. 2010). Therefore, we attempted to estimate the expression level and elucidate the tissue-specific functions of the *AsCYPs*. Grouping of the genes into different clusters on the basis of their expression pattern in various tissues including wounded Ordinary and Chi-Nan wood, leaves, flower, buds, aril, seed, stem, and callus of *A. sinensis*, indicated variations in their levels of expression among the tissues. The 21 members, namely *AsCYP71A1*, *AsCYP71A2*, *AsCYP736A1*, *AsCYP98A1*, *AsCYP89A3*, *AsCYP89A6*, *AsCYP92C1*, *AsCYP71AU2*, *AaCYP84A2*, *AsCYP71D12*, *AsCYP71D15*, *AsCYP81Q3*, *AsCYP82A9*, *AsCYP51G1*, *AsCYP89A2*, *AsCYP81Q1*, *AsCYP734A2*, *AsCYP81E3*, *AsCYP72A3*, *AsCYP84A1*, *and AsCYP71E1*, belong to 11 families and were found to be expressed in all the tissues analyzed. These genes probably involved in generation of metabolites including flavonoids, sesquiterpenoids, phenolics, sterols in all the tissues and other biological processes like BR-signaling and wound response in *A. sinensis*.

On the other hand, expression of certain members was tissue-specific, and interestingly, there were variations even within the same subfamily members. For e.g., *AsCYP71D6* was exclusively expressed in wounded Ordinary and Chi-Nan wood tissue, while *AsCYP71D20* showed expression in both seed and flower bud. Additionally, *AsCYP86A1* expressed in both leaf and flower bud, and *AsCYP75A2* exhibited expression in leaf, flower, and wounded stem. *AsCYP71D10* was found in leaf, flower, and wounded Chi-Nan wood, whereas *AsCYP82A3*, *AsCYP71D8*, *AsCYP716B2*, *AsCYP71BE4*, and *AsCYP71BE6* were solely expressed in seeds. Moreover, *AsCYP77A1* showed expression in aril, leaf, and flower bud, *AsCYP71D2* in flower and wounded stem, and *AsCYP71D3* in callus and wounded stem. *AsCYP71AU3* and *AsCYP87D1* were detected in both wounded and healthy Chi-Nan wood and stem, and *AsCYP71BE7* was found in wounded ordinary wood and stem. Additionally, *AsCYP81C1* showed higher expression specifically in leaf and callus. Information retrieved from the KEGG database revealed their tissue specific functional role. The members of the AsCYP71D and AsCYP71BE subfamilies expressed in the respective tissues are possibly involved in sesquiterpenoids biosynthesis. Similarly, members of AsCYP86A expressed in the corresponding tissues could be involved in the generation of fatty acids, whereas AsCYP75A may be related to the flavonoid biosynthesis. In the same line, members of AsCYP716B, AsCYP87D, and AsCYP82A may be involved in diterpenoid, triterpenoid biosynthesis, jasmonate and ethylene signaling, respectively. Overall, the results are consistent with other plants where difference in expression pattern within similar family members was observed indicating functional divergence. For instance, in *A. thaliana*, certain members of the CYP71B subfamily (B4, B7, and B18) showed higher expression in shoots, while other members (B6, B12, and B13) showed expression in roots. Additionally, some members (B22 and B28) were expressed in both roots and shoots (Xu et al. 2001). Similarly, members of the CYP81 family in *A. thaliana* also displayed varied expression patterns, with *CYP771A6* being specific to flowers, *CYP708A2* to roots, and *CYP83B1* and *CYP81F4* to vegetative tissues (Xu et al. 2001). In *V. vinifera*, two members, *VvCYP81D5* and *VvCYP74B2*, consistently showed high expression in all tissues studied, while *VvCYP74A1a* was expressed in ripe berries and leaves (Jiu et al. 2020). Moreover, *VvCYP71B10* elevated in berry flesh, leaves, and seeds, while *VvCYP71B24* in buds and stems. On the other hand, in rice, a significant portion (77%) of the CYPs displayed tissue-specific expression, such as *OsCYP71X14*, *OsCYP86A11*, and *OsCYP710A5*, which were specific to the endosperm, *OsCYP77A18* expressed in pistil, anther, and inflorescence, and *OsCYP96B5* found in leaves and stems (Wei at al. 2018).

The unique fragrance and industrial value of agarwood are attributed to its sesquiterpenoids profile (Tan et al. 2019). The heavily oxygenated sesquiterpenes in agarwood identified through chemical profiling were probably products of sesquiterpene oxidases of *Aquilaria* plants (Li et al. 2021, Liu et al. 2017). Previously in *A. agallocha*, we have validated the expression of certain CYPs (*AaCYP71D1*, *AaCYP71D4*, *AaCYP71D7*, *AaCYP71D10*, and *AaCYP71D11*, *AaCYP71D17*) in infected wood tissue and induced callus and demonstrated their association with sesquiterpenoids biosynthesis (Das et al. 2021). However, a tissue-wide expression study of the *Aquilaria CYPs* was lacking. This study additionally revealed the functional divergence among the 30 genes which were associated with sesquiterpenoids biosynthesis. A few (*AsCYP71D12*, *AsCYP71D15*) were expressed in all tissues, while expression of some (*AsCYP71D6*, *AsCYP71D20*, *AsCYP71D10*, *AsCYP71D8*, *AsCYP71BE4*) were restricted to 2 to 3 tissues. Notably, the higher expression of *AsCYP71D6* exclusively in wounded wood of *A. sinensis* suggests its role in the generation of sesquiterpenoids documented in agarwood (Li et al. 2021). Considering the phenomenon of agarwood formation in *Aquilaria* plants, it is plausible that CYPs respond to wounding or injury by increasing the biosynthesis of secondary metabolites, particularly sesquiterpenoids. It is concordance with the fact that wounding can potentially induce the sesquiterpene biosynthesis genes in *A. sinensis* (Sun et al. 2019). Moreover, the expression of three members, *AsCYP71D8*, *AsCYP71BE4* and *AsCYP71BE6*, exclusively in seeds might indicate their involvement in the biosynthesis of volatile sesquiterpenes, which might act as attractants for insects to facilitate rapid seed dispersal and facilitate defense against herbivores (Qin et al. 2022, Chadwick et al. 2013). Similarly, members that are particularly expressed in flower, bud, callus, and wounded stem may be involved in the generation of tissue-specific sesquiterpenoids. Overall, the evolutionary study coupled with the tissue-wide expression analysis of *AsCYPs* strongly suggests that these genes may have evolved and expanded in the *A. sinensis* genome to acquire functional divergence, in order to fulfill tissue-specific roles and adapt to the specific physiological and environmental conditions of each tissue type.

### 4.4. Enzyme Promiscuity of Sesquiterpene Oxidases in wonded wood of *Aquilaria sinensis*

Sesquiterpene oxidases (SO) exhibit a promiscuous nature, allowing them to have a versatile substrate range and produce various derivatives in plants (Hamberger et al. 2013). In our study, we identified a similar nature in AsCYP71D6 through molecular docking, demonstrating stable binding affinity towards various classes of sesquiterpenoids found in agarwood, such as eremophilane, eudesmane, and guaiane (Li et al. 2021). Moreover, the higher expression of *AsCYP71D6* specifically in wounded *A. sinensis* trees suggests its significant role in generating sesquiterpenoids derivatives during wounding and possibly in agarwood formation. Promiscuity in SO has also been reported in other plant species. For example, germacrene A oxidases (GAOs), apart from catalyzing germacrene A, also accepted germacrene D and amorphadiene as substrates in *Cynara cardunculus* (Eljounaidi et al. 2014). Similarly, members of the CYP71D family in *Nicotiana tabacum* and *Mentha piperita* can accept multiple terpenes as substrates (Wang et al. 2003, Wust et al. 2001). The enzyme promiscuity of AsCYPs potentially enhances *Aquilaria* plants’ ability to respond to wounding and infection, as it facilitates the biosynthesis of diverse oxygenated sesquiterpenoids. This feature may provide an evolutionary advantage, allowing *A. sinensis* to produce a wide range of chemical compounds that may contribute to the unique and long-lasting aroma of agarwood.

## 5. Conclusion

This study sheds light on the evolution, expansion, and tissue-wide expression of Cytochrome P450 superfamily in *Aquilaria sinensis*. The 179 full-length AsCYPs classified into 8 clans and 42 families were distributed unevenly along 8 chromosomes. Gene duplication events of the *AsCYPs* in *A. sinensis*, probably started 20 million years after diversification from *A. thaliana*, reveal that the primary driving force and mechanism responsible for the evolution and proliferation of the CYP superfamily was segmental duplication coupled with negative selection. Pathway mapping and tissue-wide expression analysis indicate that, functionally certain members of the superfamily diversified to generate secondary metabolites in a tissue specific manner. Furthermore, molecular docking analysis shows the possibilities of sesquiterpene oxidases involvement in catalyzing multiple substrates. Taken together, the functional divergence and versatile substrate selectivity collectively provide evolutionary advantage to the AsCYP superfamily. Research aimed at functional characterization of specific AsCYPs and exploring their enzyme promiscuity will provide valuable insights into the underlying mechanisms responsible for the evolution of the phytochemical profile and unique fragrance of agarwood.

## Credit authorship contribution statement

Ankur Das conceptualized, prepared the original draft, analysed the data and wrote the manuscript. Khaleda Begum validated and worked on the software and servers. Raja Ahmed worked on the software and curated data. Suraiya Akhtar helped in data curation and writing. Dr. Sofia Banu supervised, investigated and edited the manuscript.

## Acknowledgments

The authors thank Ding and team for depositing the genomic and associated data in the public database. The authors are indebted to Gauhati University for providing the technical facility, and also acknowledge the Department of Biotechnology (DBT), Government of India for providing the financial aid.

## Supplementary files

**Table S1** Full-Length 179 AsCYPs in *Aquilaria sinensis* and characteristics of the deduced proteins.

**Table S2:** The main conserved domains viz., heme-binding (red), K-helix (sky-blue), PxRx (green), and I-helix (navy blue) detected in the protein sequences of 179 AsCYPs through meme tool.

**Table S3:** Classification of the 179 AsCYPs into 8 clan and 42 families with distribution of number of genes in each family.

**Table S4:** Segmental and tandem duplicated AsCYP gene pairs with their genome location and physical distance.

**Table S5:** Orthologous gene pairs of *A. sinensis* and *A. thaliana* identified through collinearity and syntenic analysis

**Table S6:** Ka, Ks, Ka/Ks, and divergent time (Mya) of the paralogous and orthologues gene pairs.

**Table S7:** Function positions of 179 AsCYPs summarized from Uniprot, KEGG, GO and InterPro annotation.

**Table S8:** Normalized expression values of AsCYPs in different tissues in terms of log2(TPM+1).

