## Supplementary material for "Genome and transcriptome mining revealed evolutionary insights and tissue-specific expression patterns of Cytochrome P450 superfamily in *Aquilaria sinensis*": Table S2

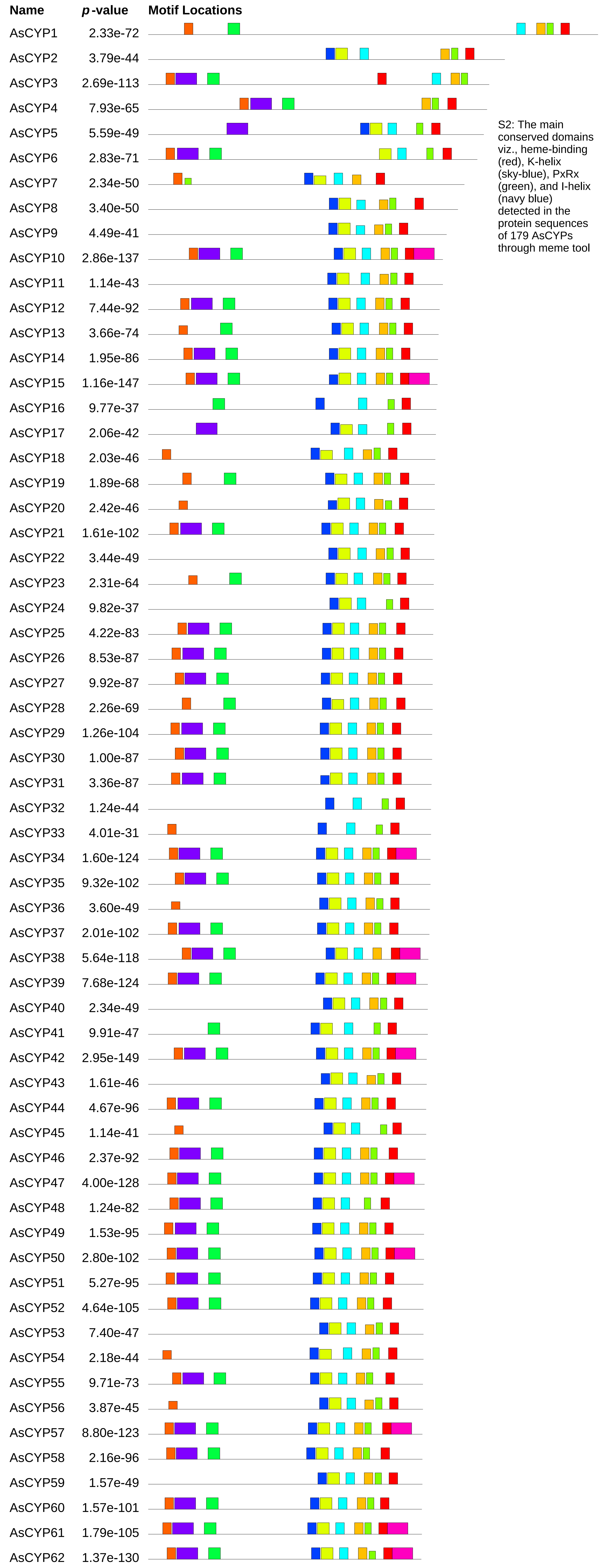


S2: The main conserved domains viz., heme-binding (red), K-helix (sky-blue), PxRx (green), and I-helix (navy blue) detected in the protein sequences of 179 AsCYPs through meme tool

| Motif | Symbol | Motif Consensus |
|---|---|---|
| 1. | | LPFGAGRRICPGISLA (Heme-binding) |
| 2. | | YLQAVIKETLRLHPPA (K-helix) |
| 3. | | WEDPEEFKPERF (PxRx motif) |
| 4. | | LPHRTLADLAKKYGPIFHLRLGSVPTVVVSSPELAKEVL |
| 5. | | PAGTRVLVNAWAIGRD |
| 6. | | YBQTDJAFAPYGPYWRQLRKJC |
| 7. | | ILAGTDTTATTLEWAM (I-helix) |
| 8. | | LANVELLLANLLYHFDWKLPBGMKPEDLDMTEKFGLTV |
| 9. | | PPGPPGLPIIGNLHLL |
| 10. | | LLKNPEVLEKAQEEVRAVVGKK |

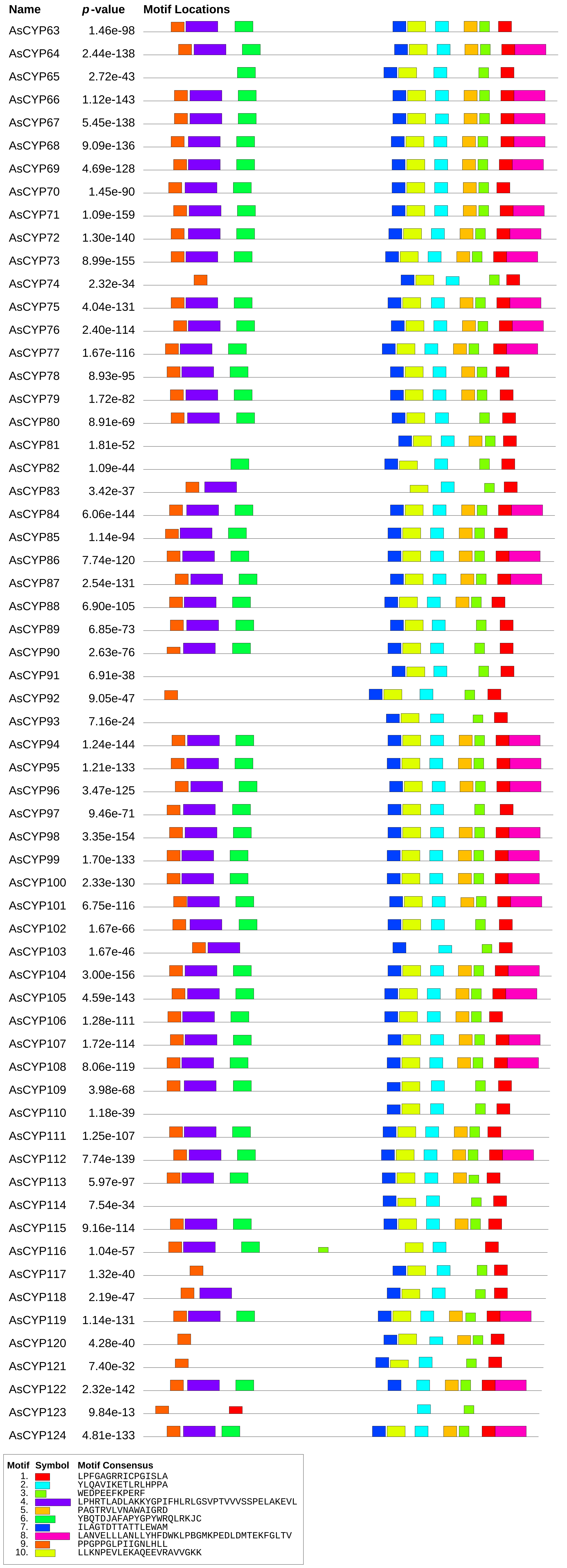
Name
p-value
Motif Locations
AsCYP63 1.46e-98
AsCYP64 2.44e-138
AsCYP65 2.72e-43
AsCYP66 1.12e-143
AsCYP67 5.45e-138
AsCYP68 9.09e-136
AsCYP69 4.69e-128
AsCYP70 1.45e-90
AsCYP71 1.09e-159
AsCYP72 1.30e-140
AsCYP73 8.99e-155
AsCYP74 2.32e-34
AsCYP75 4.04e-131
AsCYP76 2.40e-114
AsCYP77 1.67e-116
AsCYP78 8.93e-95
AsCYP79 1.72e-82
AsCYP80 8.91e-69
AsCYP81 1.81e-52
AsCYP82 1.09e-44
AsCYP83 3.42e-37
AsCYP84 6.06e-144
AsCYP85 1.14e-94
AsCYP86 7.74e-120
AsCYP87 2.54e-131
AsCYP88 6.90e-105
AsCYP89 6.85e-73
AsCYP90 2.63e-76
AsCYP91 6.91e-38
AsCYP92 9.05e-47
AsCYP93 7.16e-24
AsCYP94 1.24e-144
AsCYP95 1.21e-133
AsCYP96 3.47e-125
AsCYP97 9.46e-71
AsCYP98 3.35e-154
AsCYP99 1.70e-133
AsCYP100 2.33e-130
AsCYP101 6.75e-116
AsCYP102 1.67e-66
AsCYP103 1.67e-46
AsCYP104 3.00e-156
AsCYP105 4.59e-143
AsCYP106 1.28e-111
AsCYP107 1.72e-114
AsCYP108 8.06e-119
AsCYP109 3.98e-68
AsCYP110 1.18e-39
AsCYP111 1.25e-107
AsCYP112 7.74e-139
AsCYP113 5.97e-97
AsCYP114 7.54e-34
AsCYP115 9.16e-114
AsCYP116 1.04e-57
AsCYP117 1.32e-40
AsCYP118 2.19e-47
AsCYP119 1.14e-131
AsCYP120 4.28e-40
AsCYP121 7.40e-32
AsCYP122 2.32e-142
AsCYP123 9.84e-13
AsCYP124 4.81e-133
Motif Symbol Motif Consensus
1. LPFGAGRRICPGISLA
2. YLQAVIKETLRLHPPA
3. WEDPEEFKPERF
4. LPHRTLADLAKKYGPIFHLRLGSVPTVVVSSPELAKEVL
5. PAGTRVLVNAWAIGRD
6. YBQTDJAFAPYGPYWRQLRKJC
7. ILAGTDTTATTLEWAM
8. LANVELLLANLLYHFDWKLPBGMKPEDLDMTEKFGLTV
9. PPGPPGLPIIGNLHLL
10. LLKNPEVLEKAQEEVRAVVGKK

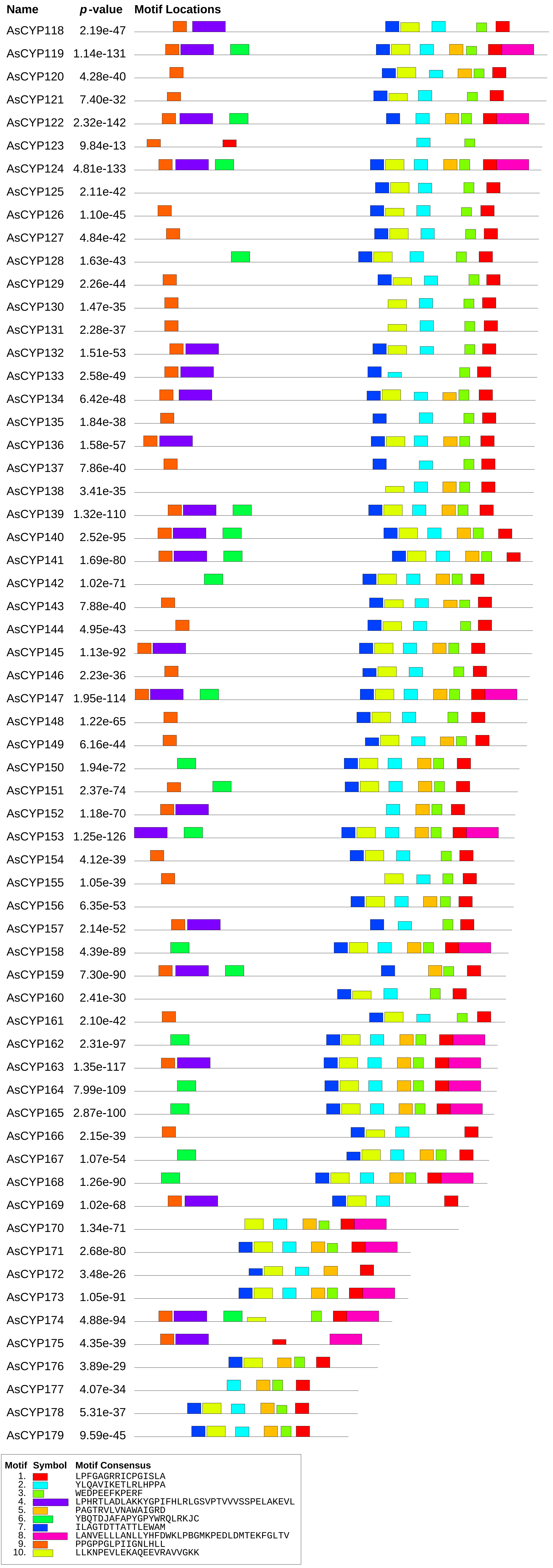
Name
p-value
Motif Locations
AsCYP118 2.19e-47
AsCYP119 1.14e-131
AsCYP120 4.28e-40
AsCYP121 7.40e-32
AsCYP122 2.32e-142
AsCYP123 9.84e-13
AsCYP124 4.81e-133
AsCYP125 2.11e-42
AsCYP126 1.10e-45
AsCYP127 4.84e-42
AsCYP128 1.63e-43
AsCYP129 2.26e-44
AsCYP130 1.47e-35
AsCYP131 2.28e-37
AsCYP132 1.51e-53
AsCYP133 2.58e-49
AsCYP134 6.42e-48
AsCYP135 1.84e-38
AsCYP136 1.58e-57
AsCYP137 7.86e-40
AsCYP138 3.41e-35
AsCYP139 1.32e-110
AsCYP140 2.52e-95
AsCYP141 1.69e-80
AsCYP142 1.02e-71
AsCYP143 7.88e-40
AsCYP144 4.95e-43
AsCYP145 1.13e-92
AsCYP146 2.23e-36
AsCYP147 1.95e-114
AsCYP148 1.22e-65
AsCYP149 6.16e-44
AsCYP150 1.94e-72
AsCYP151 2.37e-74
AsCYP152 1.18e-70
AsCYP153 1.25e-126
AsCYP154 4.12e-39
AsCYP155 1.05e-39
AsCYP156 6.35e-53
AsCYP157 2.14e-52
AsCYP158 4.39e-89
AsCYP159 7.30e-90
AsCYP160 2.41e-30
AsCYP161 2.10e-42
AsCYP162 2.31e-97
AsCYP163 1.35e-117
AsCYP164 7.99e-109
AsCYP165 2.87e-100
AsCYP166 2.15e-39
AsCYP167 1.07e-54
AsCYP168 1.26e-90
AsCYP169 1.02e-68
AsCYP170 1.34e-71
AsCYP171 2.68e-80
AsCYP172 3.48e-26
AsCYP173 1.05e-91
AsCYP174 4.88e-94
AsCYP175 4.35e-39
AsCYP176 3.89e-29
AsCYP177 4.07e-34
AsCYP178 5.31e-37
AsCYP179 9.59e-45
Motif Symbol Motif Consensus
1. LPFGAGRRICPGISLA
2. YLQAVIKETLRLHPPA
3. WEDPEEFKPERF
4. LPHRTLADLAKKYGPIFHLRLGSVPTVVVSSPELAKEVL
5. PAGTRVLVNAWAIGRD
6. YBQTDJAFAPYGPYWRQLRKJC
7. ILAGTDTTATTLEWAM
8. LANVELLLANLLYHFDWKLPBGMKPEDLDMTEKFGLTV
9. PPGPPGLPIIGNLHLL
10. LLKNPEVLEKAQEEVRAVVGKK
